# An *N*-acylphosphatidylethanolamine-LRRK2 axis controls lysosomal homeostasis in Parkinson’s disease

**DOI:** 10.64898/2026.03.19.712902

**Authors:** Francesca Palese, Carmela Giachino, Sylvie Syan, Giuliana Ottonello, Giulia Sciandrone, Carolina Filipponi, Michele Lai, Andrea Armirotti, Michela Deleidi, Chiara Zurzolo

## Abstract

*N*-acylphosphatidylethanolamines (NAPEs) are atypical glycerophospholipids that accumulate in response to cellular stress, yet their roles beyond serving as precursors of fatty-acid ethanolamines (FAEs) remain largely unexplored. Here, we identify NAPEs as endogenous regulators of leucine-rich repeat kinase 2 (LRRK2), a master controller of lysosomal homeostasis and a genetic driver of Parkinson’s disease. We show that increasing NAPE synthesis or blocking their hydrolysis inhibits LRRK2 kinase activity, enhances lysosomal function, and promotes the clearance of α-synuclein aggregates. Conversely, cells in which NAPE hydrolysis is enhanced display increased LRRK2 activation and lysosomal dysfunction. Importantly, in induced pluripotent stem cell–derived dopaminergic neurons carrying the LRRK2-G2019S variant, pharmacological inhibition of NAPE-PLD – the enzyme that degrades NAPEs – restores lysosomal activity and favors the clearance of preformed α-synuclein fibrils. Together, our findings identify NAPEs as previously unrecognized lipid regulators of LRRK2 signaling and lysosomal function, revealing a metabolic axis with therapeutic potential in Parkinson’s disease.

**Teaser:** Boosting neuronal NAPEs silences LRRK2 hyperactivity and clears α-synuclein: a lipid-based strategy for Parkinson’s disease.

## Introduction

*N*-acylphopshpatidylethanolamines (NAPEs) are low-abundance glycerophospholipids present in the membranes of all eukaryotic cells (*1*). They are synthesized through the transfer of an acyl group from phosphatidylcholine (PC) to phosphatidylethanolamine (PE) by calcium-independent phospholipase A and acyltransferases (PLAATs), which maintain basal NAPE levels (*2*), or by the calcium-dependent phospholipase PLA2G4E, which is activated in a stimulus-dependent manner to markedly increase NAPE production under pathological conditions (*3*, *4*). NAPEs are hydrolyzed by the NAPE-specific phospholipase D (NAPE-PLD) (*5*, *6*), generating fatty-acid ethanolamines (FAEs) and phosphatidic acid (PA).

FAEs mediate diverse physiological and pathological processes, including neurotransmission (*7*, *8*), pain (*9*), energy homeostasis (*10*, *11*) and inflammation (*12*, *13*), through receptors such as cannabinoid receptors, TRPV1, GPR119 and PPAR-α. Consequently, NAPEs have traditionally been viewed mainly as precursors of FAEs rather than molecules with intrinsic biological activity. However, NAPE production is dynamically regulated by physiological (*7*, *14*) and pathological stimuli (*15–17*) including ischemia (*18*), excitotoxicity (*19*) and neurotoxic insults (*20–22*). Under pathological conditions, NAPE levels often exceed those of FAEs, suggesting functions independent of downstream FAE signaling. Consistent with this idea, NAPEs have been shown to stabilize cellular membranes (*23*, *24*), regulate membrane fusion (*25*) and lipid raft organization (*26*), thereby influencing protein association at the membranes (*27*, *28*). Notably, NAPEs accumulate in the brains of mouse models of Parkinson’s disease (PD) (*21*, *29*), where they exert neuroprotective effects against 6-OHDA-induced toxicity (*29*). In contrast, circulating NAPE levels are decreased in the plasma of female PD patients (*30*).

These observations suggest that NAPEs may participate in adaptive lipid response to neuronal stress in PD. However, the molecular mechanisms through which NAPEs influence neuronal survival and disease-relevant pathways remain largely unknown.

One potential link between NAPE signaling and PD pathogenesis is the regulation of membrane-associated signaling proteins that control endolysosomal trafficking. Among there, leucine-rich repeat kinase 2 (LRRK2) has emerged as a central regulator of endolysosomal dynamics and lysosomal function whose dysregulation is strongly linked to PD (*31*). Mutations in the *LRRK2* gene represent one of the most common genetic causes of PD, accounting for ∼1-2% of all cases worldwide but reaching up to 30-40% of cases in certain founder populations (*32*). Previous work demonstrated that NAPE accumulation reduces LRRK2 association with neuronal membrane fractions, whereas NAPE hydrolysis promotes its membrane recruitment (*28*). Here, we investigate the molecular mechanism underlying NAPE-mediated neuroprotection, focusing on LRRK2 regulation. We find that NAPE accumulation — either through enhanced synthesis or inhibition of their hydrolysis — reduces LRRK2 kinase activity and improves lysosomal function, thereby promoting the clearance of α-synuclein pre-formed fibrils (PFFs). Conversely, overexpression of the hydrolyzing enzyme NAPE-PLD increases LRRK2 protein abundance and kinase activity, impairs lysosomal function, and decreases α-synuclein PFF clearance.

Importantly, in induced pluripotent stem cell-derived dopaminergic neurons carrying the hyperactivating LRRK2-G2019S variant, pharmacological inhibition of NAPE-PLD restores lysosomal activity and enhances α-synuclein PFF clearance. Together, these findings identify NAPEs as previously unrecognized regulators of LRRK2 activity and lysosomal function, with direct implications for PD pathogenesis, and highlight NAPE-PLD inhibitors as promising therapeutic candidates.

## Results

### NAPEs accumulation potentiates lysosomal activity via LRRK2 inhibition

To investigate the effects of NAPE accumulation, we genetically modulated NAPE metabolism in catecholaminergic SH-SY5Y cells using two approaches: (i) overexpression of the synthesizing enzyme PLA2G4E and (ii) knockdown of the hydrolyzing enzyme NAPE-PLD (**Fig. 1A**). Using a CRISPR-Cas9 strategy we obtained a partial knockdown (KD), resulting in an approximatively 30% reduction in *NAPE-PLD* mRNA (**Fig. 1B**), and a 30-50% decrease in protein levels (depending on the technique used for the measurement, **Fig. 1C, D**). This led to a 30% increase in total NAPE levels (**Fig. 1E**). Consistent with a hypomorphic condition, no significant compensatory changes were observed in *PLA2G4E* mRNA or protein (**Fig. S1 A-C**).

**Figure 1.**
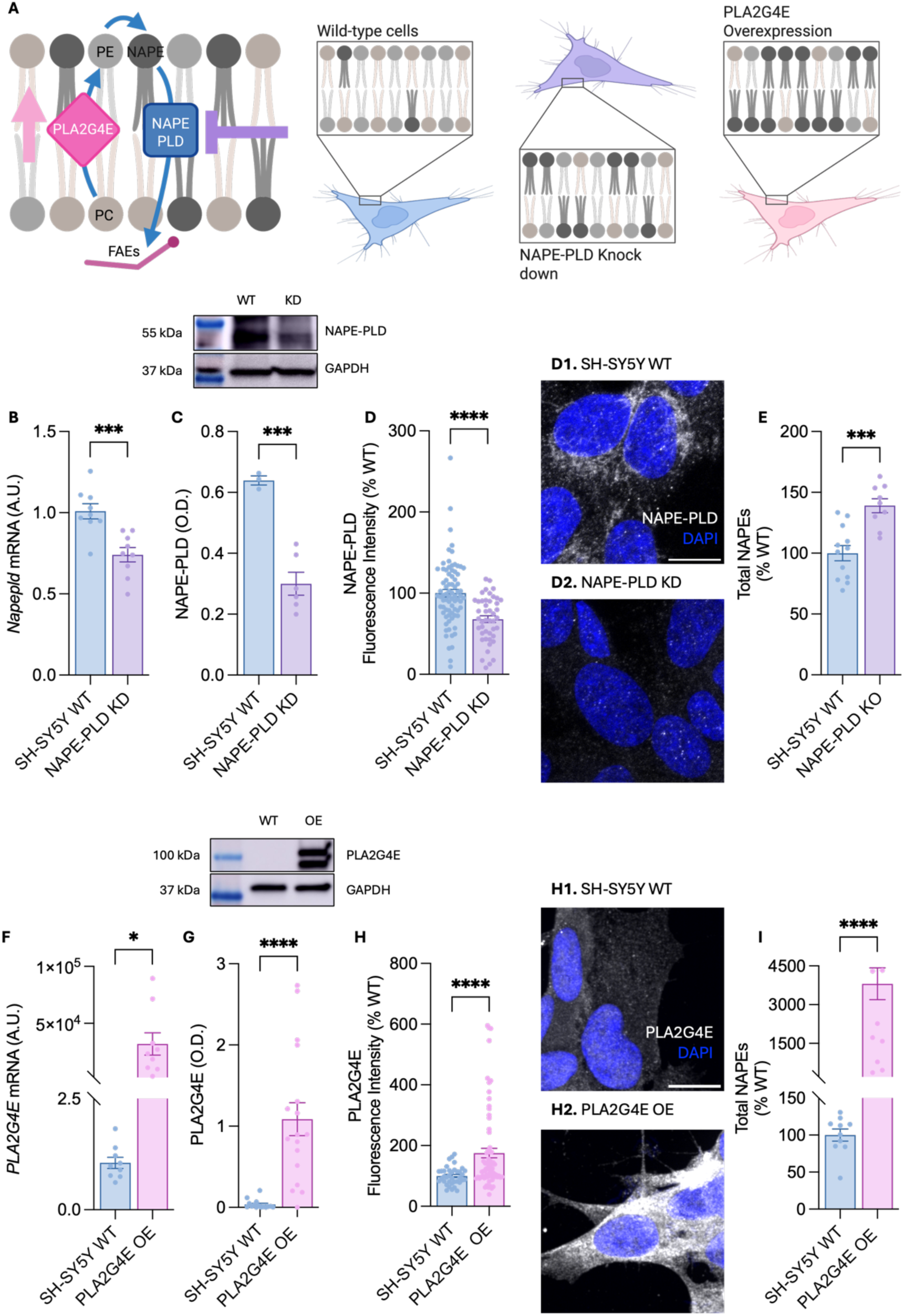
Generation of catecholaminergic SH-SY5Y neuronal cells accumulating NAPEs. (**A**) Schematic representation of the experimental strategy. Stable SH-SY5Y cell lines were generated either by knocking down NAPE-PLD using CRISP-Cas9 approach or by overexpressing PLA2G4E, both manipulations led to NAPE accumulation at the cell membranes. (**B**) *NAPE-PLD* mRNA levels in NAPE-PLD-knockdown (KD; violet bars) and wild-type (WT; blue bars) cells. (**C**) Western blot analysis of NAPE-PLD protein levels in KD and WT cells. Top, representative blot; bottom, densitometric quantification normalized to glyceraldehyde 3-phosphate dehydrogenase (GAPDH). (**D**) Immunofluorescence analysis of NAPE-PLD (gray) with representative images (**D1, D2**) and quantification. Nuclei were counterstained with DAPI. (**E**) Total NAPE levels in KD and WT cells. (**F**) *PLA2G4E* mRNA levels in PLA2G4E-overexpressing (OE; pink bars) and WT (blue bars) cells. (**G**) Western blot analysis of PLA2G4E protein levels in OE and WT cells. Top, representative blot; bottom, densitometric quantification normalized to GAPDH. (**H**) Immunofluorescence analysis of PLA2G4E (gray) with representative images (**H1, H2**) and quantification. Nuclei were counterstained with DAPI. (**I**) Total NAPE levels in OE and WT cells. Error bars represent SEM. *P < 0.05, **P < 0.01 ***P < 0.001, ****P < 0.0001 Student’s *t*-test with Welch’s correction. Scale bars, 10 μm.

In parallel, SH-SY5Y cells overexpressing (OE) the synthesizing enzyme PLA2G4E (*33*) displayed a massive upregulation of both mRNA (∼3×10^4^-fold) and protein (∼30-fold) levels (**Fig 1F-H**). In line with a strong potentiation of the synthetic pathway, these cells exhibited a 70-fold increase in total NAPE content (**Fig. 1 I**). Notably, NAPE-PLD protein levels were downregulated in these conditions (**Fig. S1 E, F**), likely reflecting an adaptive response aimed at limiting excessive FAE production.

Previous studies in NAPE-PLD knockout mouse brains and in NAPE-PLD silenced neuronal cells showed a reduced LRRK2 fraction associated with membranes (*28*); however, the underlying mechanisms and functional consequences of this observation were not investigated. Here, we show that both NAPE-PLD KD and PLA2G4E OE reduce LRRK2 abundance in neuronal cells (**Fig. 2 A**). Moreover, the LRRK2 kinase activity measured by western blot using phosphorylation of Rab8a at T72 — a LRRK2-specific phosphorylation site — was lowered in both NAPE-PLD KD and PLA2G4E OE cells compared with WT SH-SY5Y cells (**Fig. 2 C**). Furthermore, we observed diminished LRRK2 localization at lysosomes (**Fig. 2B, D**) in both NAPE-PLD KD and PLA2G4E OE cells relative to WT cells. Given that LRRK2 can be recruited to lysosomes in response to lysosomal stress or damage (*34*, *35*) this is consistent with improved lysosomal health under conditions of NAPE accumulation.

**Figure 2.**
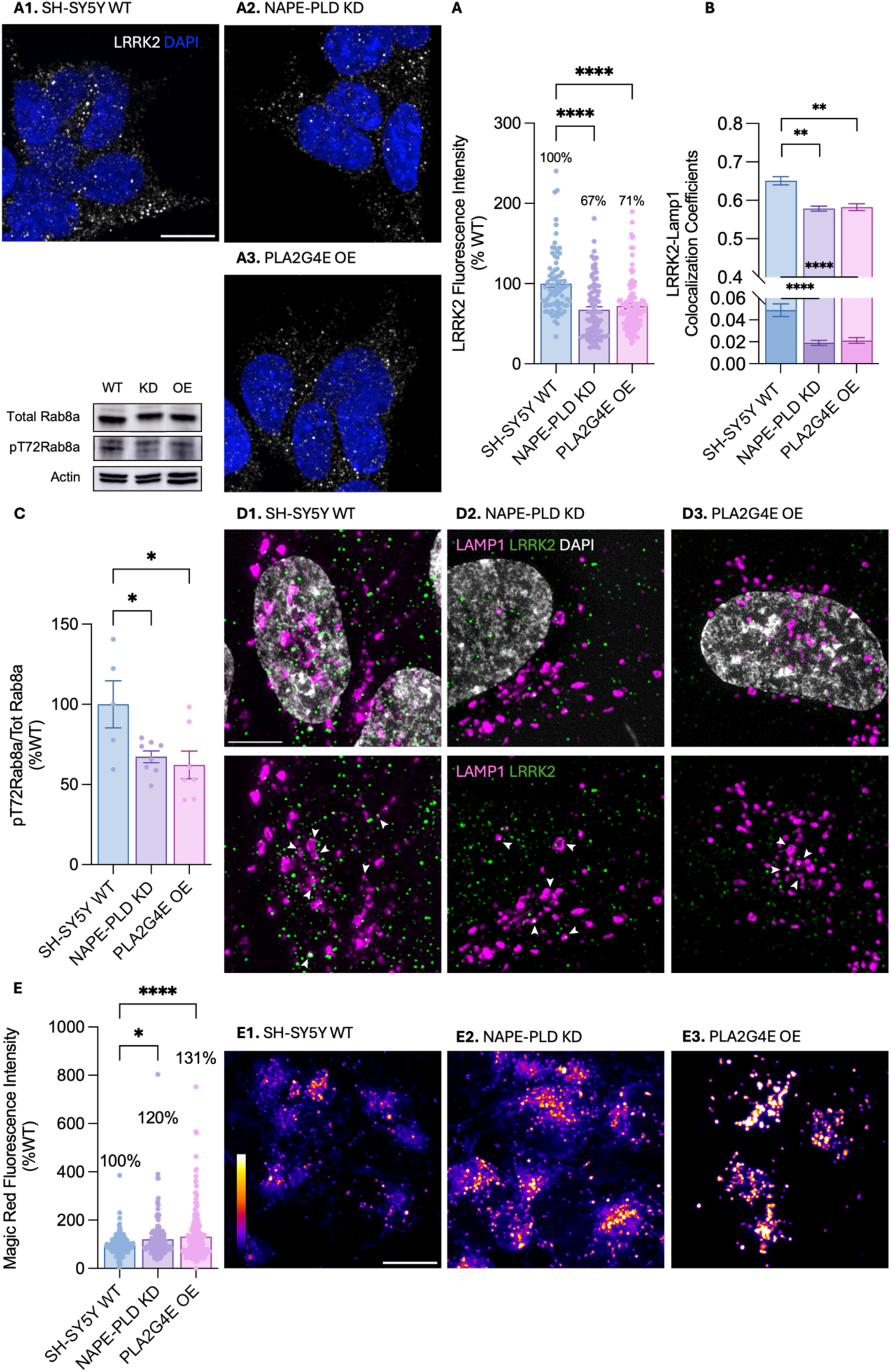
Neuronal cells accumulating NAPEs downregulate LRRK2 protein and potentiate lysosomal activity. (**A**) Immunofluorescence analysis of LRRK2 (gray) with representative images (**A1–A3**) and quantification in wild-type (WT; **A1**, blue), NAPE-PLD knockdown (KD; **A2**, violet), and PLA2G4E-overexpressing (OE; **A3**, pink) SH-SY5Y cells. Nuclei were counterstained with DAPI. (**B**) Quantification of colocalization between LAMP1-positive structures and LRRK2, expressed as Pearson’s correlation coefficient (upper bars) and Manders’ overlap coefficient (lower bars) for the fraction of LAMP1 signal overlapping with LRRK2. (**C**) Western blot analysis of Rab8a and phosphorylated Rab8a (pT72-Rab8a) in KD and OE compared with WT cells. Top, representative blot; bottom, densitometric quantification normalized to actin. (**D**) Representative super-resolution immunofluorescence images of LAMP1 (magenta) and LRRK2 (green) in WT (**D1**), NAPE-PLD KD (**D2**), and PLA2G4E OE (**D3**) cells. Nuclei were stained with DAPI (gray). Scale bar, 5 μm. (**E**) Fluorescence intensity quantification and representative images of the Magic Red–cathepsin B assay, indicating lysosomal degradative capacity per cell, in WT (**E1**), NAPE-PLD KD (**E2**), and PLA2G4E OE (**E3**) cells. Representative images are displayed using the FIRE lookup table (LUT) to indicate fluorescence intensity. Scale bars, 10 μm. Mean values are indicated in the graphs; error bars represent SEM. *P < 0.05, ****P < 0.0001; one-way ANOVA with Tukey’s multiple-comparison test.

LRRK2 is an established regulator of lysosomal degradative activity. LRRK2 knockout or kinase inhibition has been shown to increase lysosomal proteolytic activity, whereas pathogenic, hyperactive LRRK2 mutations associated with PD impair lysosomal degradative activity (*36–38*). In accordance with an inhibitory function of LRRK2 on lysosomal activity, we observed increased lysosomal cathepsin B activity in cells where NAPEs accumulate and LRRK2 is decreased in a NAPE-dose-dependent manner (+20% in NAPE-PLD KD cells and + 31% in PLA2G4E OE cells; **Fig. 2E**). Interestingly, treatment with the calcium ionophore ionomycin - which is reported to stimulate the production of NAPEs (*39*)(**Fig. S2A**)- further improved lysosomal activity in both NAPE-enriched cell lines (**Fig. S2B**).

Since increased lysosomal enzyme activity can also occur as a compensatory response to impaired autophagosome-lysosome fusion, we assessed autophagic flux to avoid misinterpretation. Cells were treated with the autophagosome-lysosome fusion inhibitor Bafilomycin A1, and p62-Lamp1 colocalization was measured alongside LC3 immunoblot analysis. Under basal conditions, no significant differences were observed between cell lines (**Fig. S3A**). As expected, Bafilomycin A1 treatment led to the accumulation of p62-Lamp1-positive structures across all conditions (**Fig. S3A**), accompanied by increased LC3-II levels (**Fig. S3B**), consistent with intact autophagosome-lysosome fusion. Together, these data indicate that NAPE accumulation enhances lysosomal activity rather than reflecting impaired autophagic flux.

Furthermore, no differences were observed in TFEB nuclear localization or total levels across cell lines (**Fig. S3C-E**), nor in the transcription of TFEB target genes (**Fig. S3 F-I**), indicating that NAPE-mediated lysosomal protection occurs independently of TFEB activation, the master regulator of lysosomal biogenesis (*40*).

### NAPEs-enriched cells clear exogenous pre-formed α-synuclein fibrils more efficiently

We next investigated the effect of NAPE accumulation under pathological conditions, specifically following exposure to exogenous pre-formed *α*-synuclein (*α*-syn) fibrils (PFFs). *α*-Syn PFFs are well known to induce lysosomal stress and activate LRRK2 kinase activity in neurons (*41*, *42*), thereby establishing a negative feedback loop on lysosomal activity. Interestingly, following overnight exposure of WT, NAPE-PLD KD and PLA2G4E OE cells to *α*-syn PFFs we detected a ∼2-fold increase in LRRK2 protein levels in WT cells, whereas the increase was very modest in NAPE-enriched cells (+4% and +34% in NAPE-PLD KD and PLA2G4E OE cells, respectively; **Fig. 3 A**). These data support NAPE-mediated negative regulation of LRRK2 protein levels, in addition to reduced LRRK2 recruitment to lysosomes (**Fig. 3 B**). Furthermore, consistent with their enhanced lysosomal activity, both NAPE-PLD KD and PLA2G4E OE cells showed fewer *α*-syn aggregates after overnight PFF treatment, followed by a 6-h washout period, compared with WT SH-SY5Y cells (**Fig. 3 C**).

**Figure 3.**
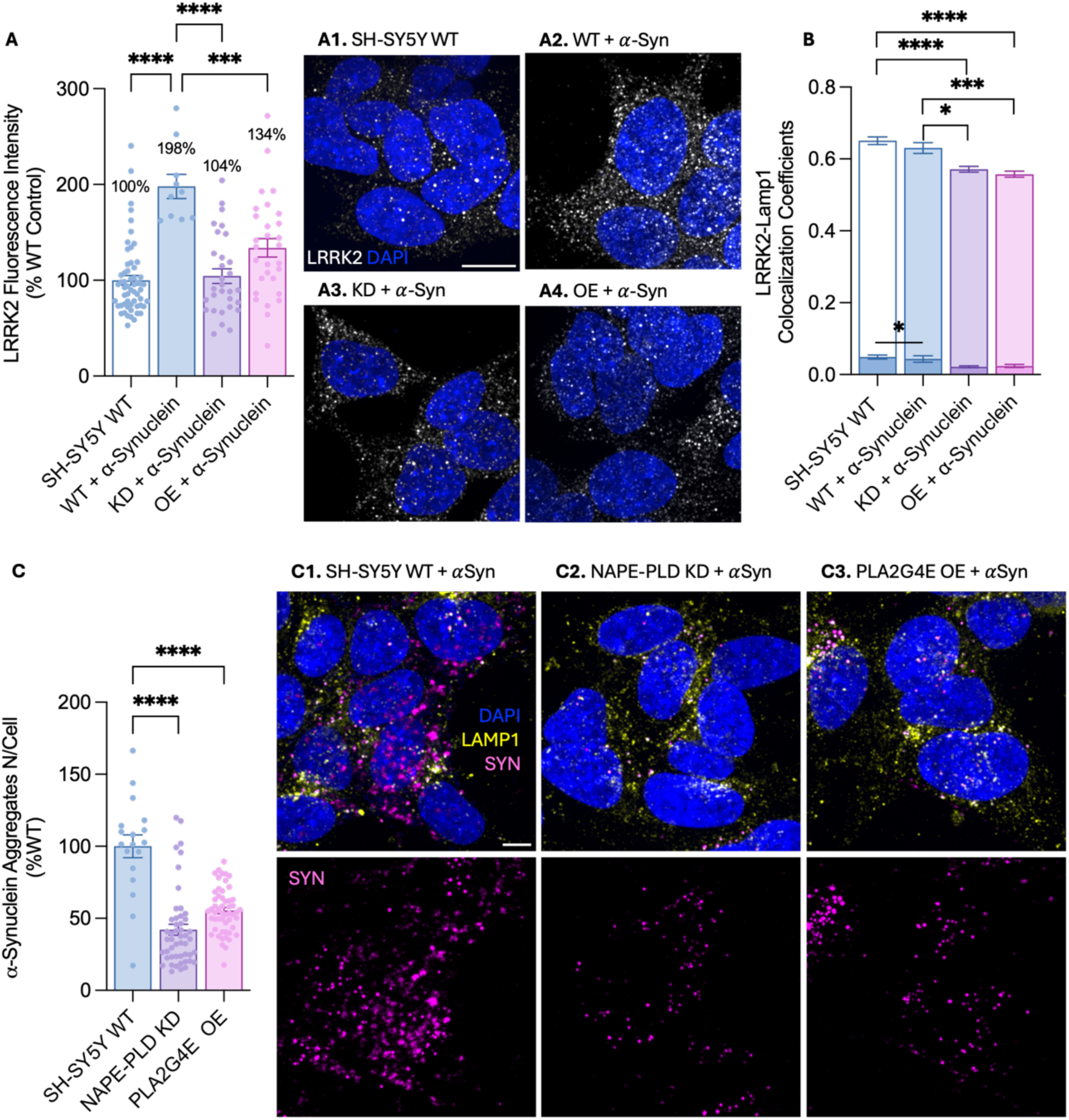
Neuronal cells accumulating NAPEs prevent LRRK2 accumulation following α-synuclein treatment and enhance aggregate clearance. (**A**) Immunofluorescence analysis of LRRK2 (gray) with representative images (**A1-A4**) and quantification in WT (**A1, A2**), NAPE-PLD-knockdown (KD; **A3**), and PLA2G4E-overexpressing (OE; **A4**) cells treated or not with α-syn (**A2-A4**). (**B**) Quantification of the colocalization between LAMP1-positive structures and LRRK2, expressed as Pearson’s coefficient (upper bars) and Manders’ overlap coefficient (lower bars) for the fraction of LAMP1 signal overlapping with LRRK2. (**C**) Quantification and representative images of α-syn aggregate number per cell in WT (**C1**), NAPE-PLD KD (**C2**) and PLA2G4E OE (**C3**) cells. LAMP1 is shown in yellow, α-syn in pink, and nuclei were counterstained with DAPI (blue). Scale bar, 10 μm. Mean values are indicated in the graphs; error bars represent SEM. ***P < 0.001, ****P < 0.0001; one-way ANOVA with Tuckey’s multiple-comparison test.

### Lysosomal activity and α-synuclein fibrils clearance are impaired in NAPE-PLD-overexpressing cells

To strengthen these findings, we next generated SH-SY5Y cells overexpressing NAPE-PLD. In these cells, *NAPE-PLD* mRNA and protein levels increased by approximately 10^3^-fold and 5-fold respectively (**Fig. 4 A-D**). This overexpressing markedly enhance NAPE hydrolysis at cellular membranes under basal condition and promote FAEs production (*43*), resulting in a ∼40% decrease in NAPE content (**Fig. 4 E**). Consistent with the non-rate limiting role of NAPE-PLD in FAE biosynthesis (*44*), no major compensatory effects were detected in PLA2G4E expression levels (**Fig. S4 A-C**).

**Figure 4.**
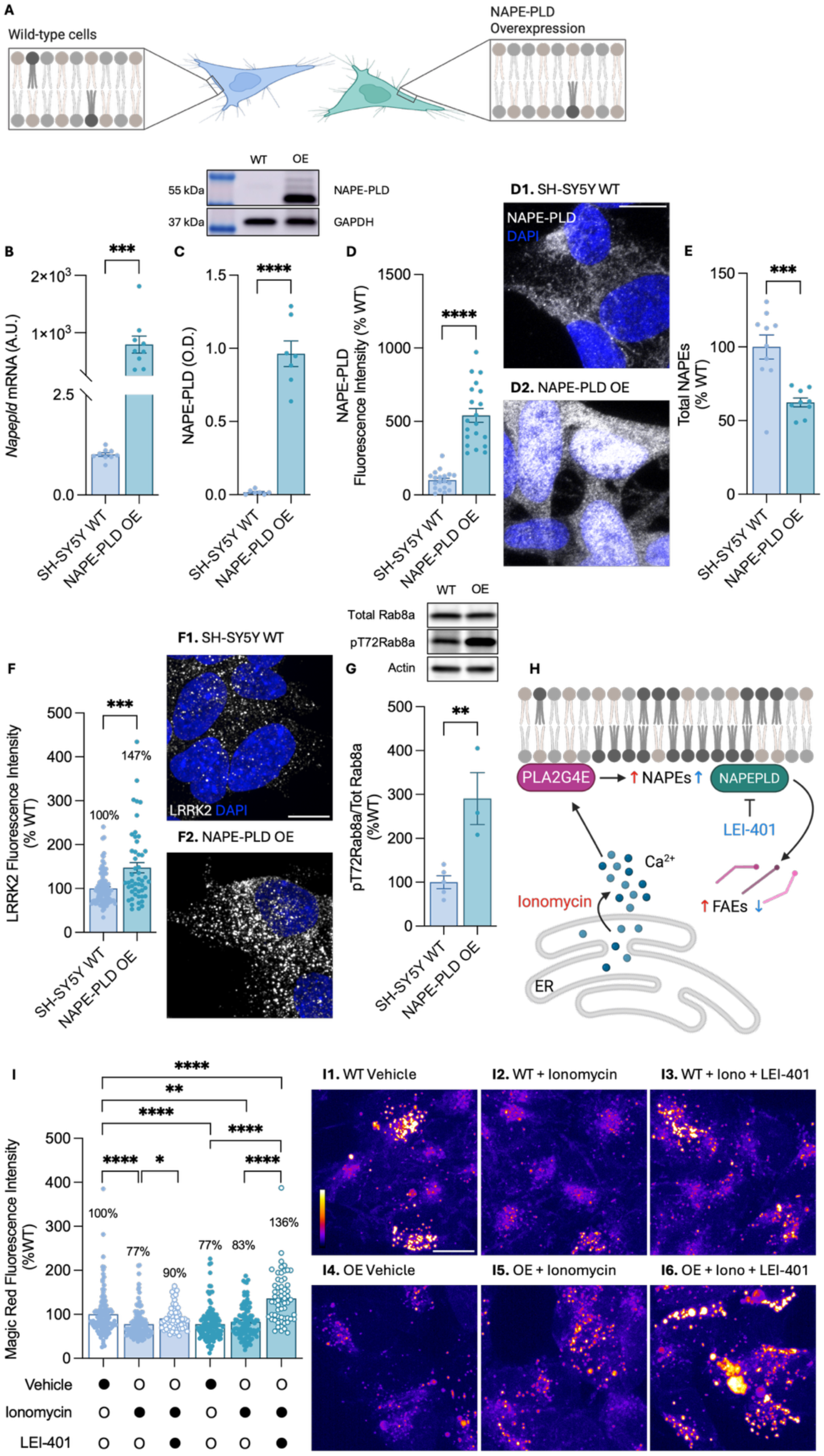
NAPE-PLD overexpression upregulates LRRK2 protein and impairs lysosomal activity. (**A**) Schematic representation of the generation of stable SH-SY5Y cell lines overexpressing NAPE-PLD, leading to reduced NAPE levels at cellular membranes. (**B**) *NAPE-PLD* mRNA levels in NAPE-PLD-overexpressing (OE; green) and wild-type (WT; blue) cells. (**C**) Western blot analysis of NAPE-PLD protein levels in OE and WT cells. Top, representative blot; bottom, densitometric quantification normalized to GAPDH. (**D**) Immunofluorescence analysis of NAPE-PLD (gray) with representative images (**D1, D2**) and quantification. (**E**) Total NAPE levels in OE and WT cells. (**F**) Immunofluorescence analysis of LRRK2 (gray) with representative images (**F1, F2**) and quantification in WT (**F1**) and NAPE-PLD OE (**F2**) cells. Nuclei were counterstained with DAPI (blue). (**G**) Western blot analysis of Rab8a and phosphorylated Rab8a (pT72-Rab8a) in NAPE-PLD OE and WT cells. Top: representative blot; bottom, densitometric quantification normalized to actin. (**H**) Schematic representation of the experimental strategy. Ionomycin treatment induces intracellular Ca^2+^ release, promoting NAPE production and degradation in NAPE-PLD OE cells. NAPE-PLD activity was inhibited by overnight treatment with the NAPE-PLD inhibitor LEI-401. Ionomycin was administered together with Magic Red reagent for 30 min before image acquisition. (**I**) Fluorescence intensity quantification and representative images of the Magic Red – cathepsin B assay, indicating lysosomal degradative capacity per cell, in untreated WT (**I1,** white bar and blue dots) and NAPE-PLD OE cells (**I4,** white bar and green dots), ionomycin-treated WT (**I2,** blue bar and blue dots) and NAPE-PLD OE cells (**I5,** green bar and green dots) or WT (**I3,** blue bar white dots) and NAPE-PLD OE cells (**I6**, green bar and white dots) treated with ionomycin and LEI-401. Representative images are displayed using the FIRE lookup table (LUT) to indicate fluorescence intensity. Scale bars, 10 μm. Mean values are indicated in the graphs; error bars represent SEM. *P < 0.05, **P < 0.01, ***P < 0.001, ****P < 0.0001; Student’s *t*-test with Welch’s correction (**B-G**) and two-way ANOVA with Tuckey’s multiple-comparisons test (**H**).

In contrast to NAPE-enriched cells, NAPE-PLD OE cells exhibited elevated LRRK2 protein levels and kinase activity (**Fig. 4 F, G**). These cells displayed markedly reduced cathepsin B activity relative to WT (77% of WT levels; **Fig. 4I**), consistent with LRRK2’s known suppression of lysosomal hydrolases. Notably, ionomycin treatment -which boosts NAPE production and, in NAPE-PLD-overexpressing cells, is expected to promote robust FAEs generation (**Fig. 4H**)- failed to rescue cathepsin B activity (**Fig. 4I**). This suggests that lysosomal impairment is driven by altered NAPE levels rather than by FAE or phosphatidic acid byproducts. Importantly, the lysosomal impairment observed in NAPE-PLD OE cells was reversed by treatment with LEI-401 (a NAPE-PLD inhibitor) (**Fig. S5 A**). Furthermore, when LEI-401 and ionomycin treatments were combined, lysosomal activity reached 136% of WT levels, consistent with maximal NAPE accumulation (**Fig. 4I**). In line with our observation of elevated LRRK2 activity in NAPE-PLD OE cells, overnight treatment with the LRRK2 pharmacological inhibitor MLi-2, not only improved lysosomal activity in WT cells, but it also rescued the lysosomal impairment detected in NAPE-PLD OE cells (**Fig. S6**), supporting the conclusion that NAPE levels regulate lysosomal activity through modulation of LRRK2 abundance and kinase activity.

Following Bafilomycin A1 treatment, NAPE-PLD OE cells displayed reduced p62-Lamp1 colocalization compared to WT cells (**Fig. S7A**), along with a non-significant decrease in LC3-II accumulation (**Fig. S7B**), suggesting impaired autophagosome-lysosome fusion, consistent LRRK2 overexpression (*45*). Moreover, these cells exhibited increased TFEB levels in both cytosolic and nuclear compartments (**Fig. S7B-E**), accompanied by elevated transcription of *CTSD* and *ATP6V1H1* (**Fig. S7H-I**), indicative of a compensatory response aimed at restoring lysosomal homeostasis.

Finally, following overnight treatment with *α*-syn PFFs, LRRK2 protein levels increased further in NAPE-PLD OE cells (∼3-fold relative to untreated WT cells and ∼1.5-fold relative to WT cells treated with *α*-syn; **Fig. 5A**), in contrast to NAPE-enriched cells, where the *α*-syn-induced increase in LRRK2 levels was contained. Consistent with reduced lysosomal degradative capacity, the number of *α* -syn aggregates after a 6-hour washout period were higher in NAPE-PLD OE SH-SY5Y cells than in WT cells (**Fig. 5 B**). Together, these results identify a previously unrecognized role for NAPEs in regulating lysosomal activity and the handling of protein aggregates in neurons through modulation of LRRK2 activity.

**Figure 5.**
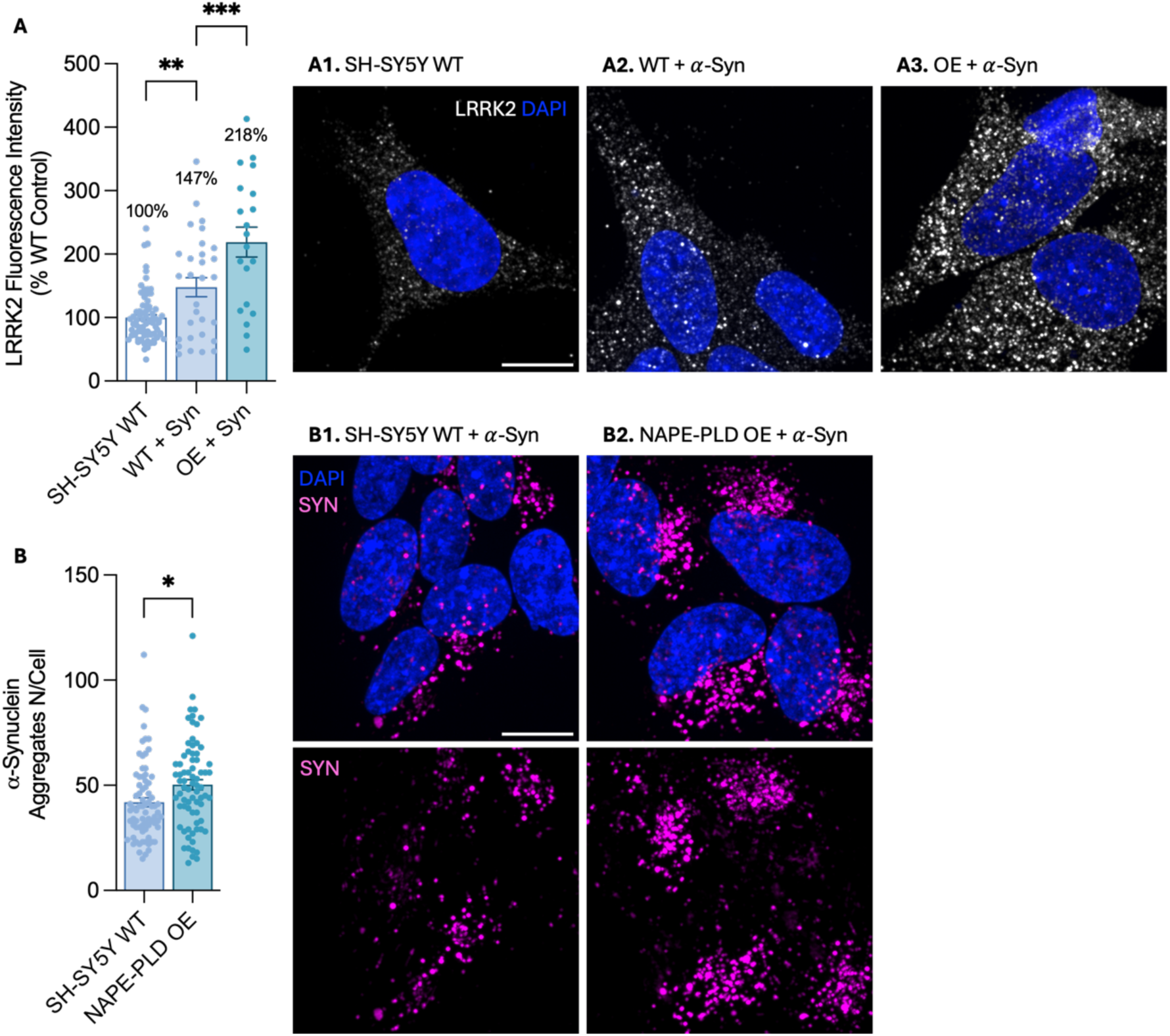
NAPE-PLD overexpression enhances LRRK2 accumulation following α-synuclein treatment and impairs aggregate clearance. (**A**) Immunofluorescence analysis of LRRK2 (gray) with representative images (**A1-A3**) and quantification in WT (**A1, A2**), and NAPE-PLD OE (**A3**) SH-SY5Y cells treated or not with α-synuclein (α-syn; **A2, A3**). (**B**) Quantification and representative images of α-syn aggregate number per cell in WT (**B1**) and NAPE-PLD OE (**B2**) cells. α-Syn is shown in pink, and nuclei were counterstained with DAPI. Scale bar, 10 μm. Mean values are indicated in the graphs; error bars represent SEM. *P<0.05, ***P < 0.001, ****P < 0.0001; one-way ANOVA with Tuckey’s multiple-comparison test (**A**) and Student’s *t*-test with Welch’s correction (**B**).

### NAPE-PLD inhibition restores lysosomal activity and α-synuclein fibril clearance in G2019S-LRRK2 dopaminergic neurons

We next evaluated the therapeutic potential of NAPE-PLD inhibition in human iPSC-derived dopaminergic neurons (hDANs) carrying the PD-linked LRRK2-G2019S mutation and their isogenic control line. This gain-of-function variant enhances LRRK2 kinase activity (*46*) and impairs lysosomal function (*47*). hDANs were analyzed at differentiation days 21-24, when they express neuronal markers (TUJ1 and MAP2) and the dopaminergic marker tyrosine hydroxylase (TH; **Fig. S8 A, B**), and display spontaneous activity appropriate membrane potential, and other neurophysiological features indicative of functional maturity (*48*). Notably, LRRK2 protein levels are comparable between control and G2019S-LRRK2 lines (*49*).

Cells were treated overnight with the NAPE-PLD inhibitor LEI-401, and lysosomal activity was assessed using the Magic Red Cathepsin B assay. Consistent with LRRK2-mediated lysosomal impairment, G2019S-LRRK2 neurons exhibited fewer active lysosomes than isogenic controls, an effect rescued by LEI-401 treatment (**Fig. 6 A, B**). These findings indicate that increasing NAPE levels through NAPE-PLD inhibition counteracts LRRK2 hyperactivation and restore lysosomal function.

**Figure 6.**
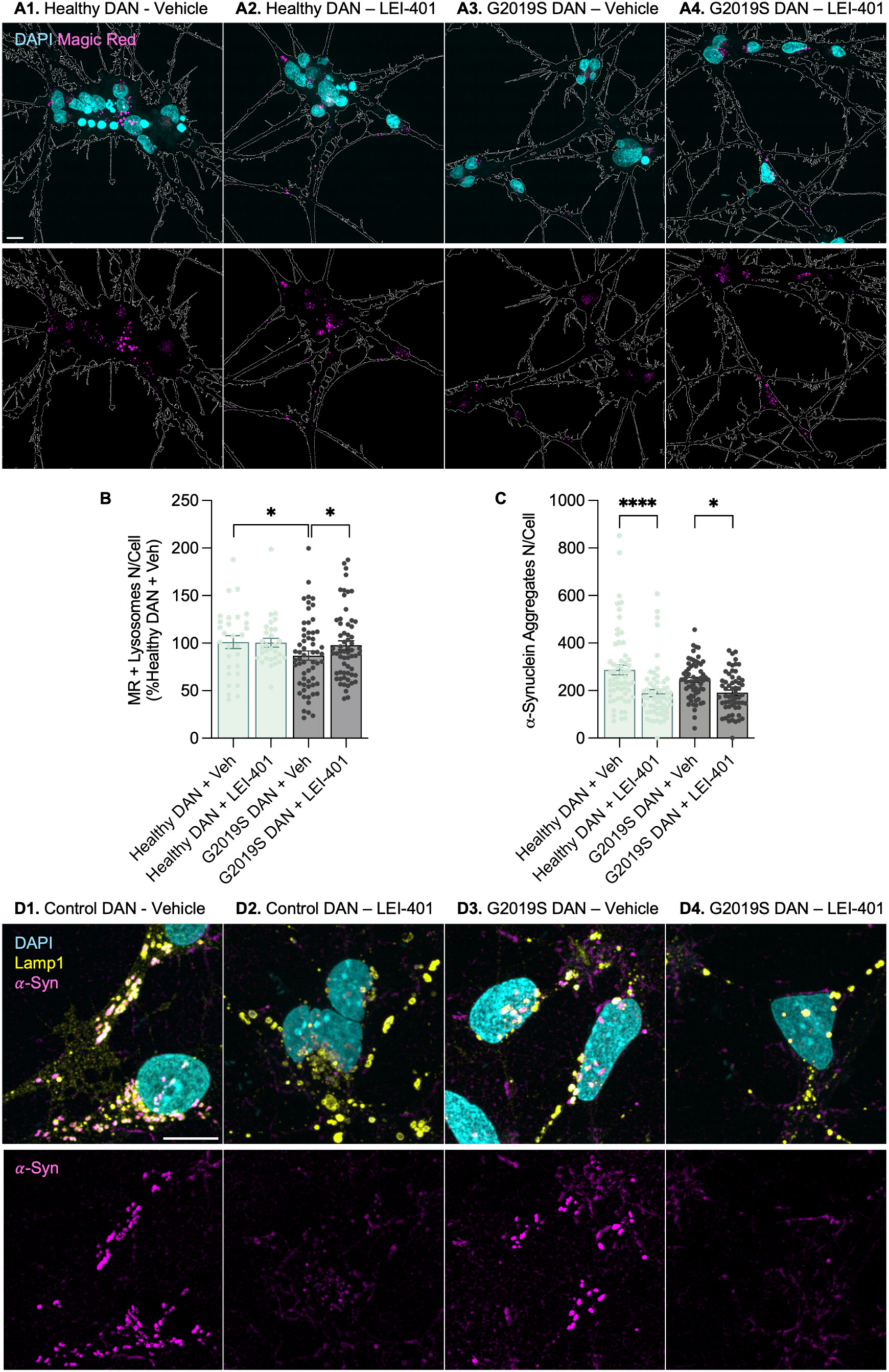
NAPE-PLD inhibition restores lysosomal activity and α-synuclein fibril clearance in G2019S-LRRK2 dopaminergic neurons. (**A**) Representative images of the Magic Red – cathepsin B assay, indicating lysosomal degradative capacity per cell, in healthy dopaminergic neurons (DANs; **A1, A2**) and G2019S-LRRK2 mutant DANs (**A3, A4**) treated (**A2, A4**) or not (**A1, A3**) with the NAPE-PLD inhibitor LEI-401. (**B**) Fluorescence intensity quantification of the Magic Red assay in healthy DANs (green) and G2019S-LRRK2 mutant DANs (gray). (**C, D**) Quantification and representative images of α-synuclein (α-syn) aggregate number per cell in healthy DANs (**D1, D2**) and G2019-LRRK2 mutant DANs (**D3, D4**) treated (**D2, D4**) or not (**D1, D3**) with LEI-401. α-Syn is shown in pink, LAMP1 yellow, and nuclei were counterstained with DAPI. Scale bars, 10 μm. *P < 0.05, ****P < 0.0001; two-way ANOVA with Tuckey’s multiple-comparison test.

We next assessed whether this rescue translated into improved handling of pathological *α*-syn. hDANs were treated with *α*-syn PFFs in the presence or absence of LEI-401. Both G2019S-LRRK2 neurons and isogenic controls efficiently internalized PFFs after 48 h incubation (**Fig. 6 C**), with comparable baseline uptake between genotypes. Strikingly, LEI-401 treatment significantly reduced the number of intracellular *α*-syn aggregates per cell in both mutant and control neurons compared to untreated conditions, demonstrating enhanced aggregate clearance. Together, these results establish NAPE-PLD inhibition as a mechanism to restore lysosomal function and promote pathological protein clearance in human dopaminergic neurons. These findings identify NAPEs as upstream modulators of LRRK2-dependent lysosomal dysfunction and highlight NAPE-PLD as a promising therapeutic target for Parkinson’s disease and related neurodegenerative disorders characterized by impaired lysosomal proteostasis.

## Discussion

In the present study, we identify NAPEs as previously unrecognized lipid regulators of LRRK2 signaling and lysosomal function in neuronal cells and human dopaminergic neurons. By genetically and pharmacologically manipulating NAPE synthesis and hydrolysis, we define a lipid-kinase axis in which NAPE accumulation (i) restrains LRRK2 kinase activity, (ii) preserves lysosomal degradative capacity, and (iii) promotes *α*-synuclein aggregate clearance, whereas NAPE depletion has the opposite effect (resumed in **Fig. 7**). These findings provide a mechanistic framework linking stress-induced NAPE production to endolysosomal homeostasis and suggest that NAPE-PLD is a viable target to counteract LRRK2 hyperactivation in PD.

**Figure 7.**
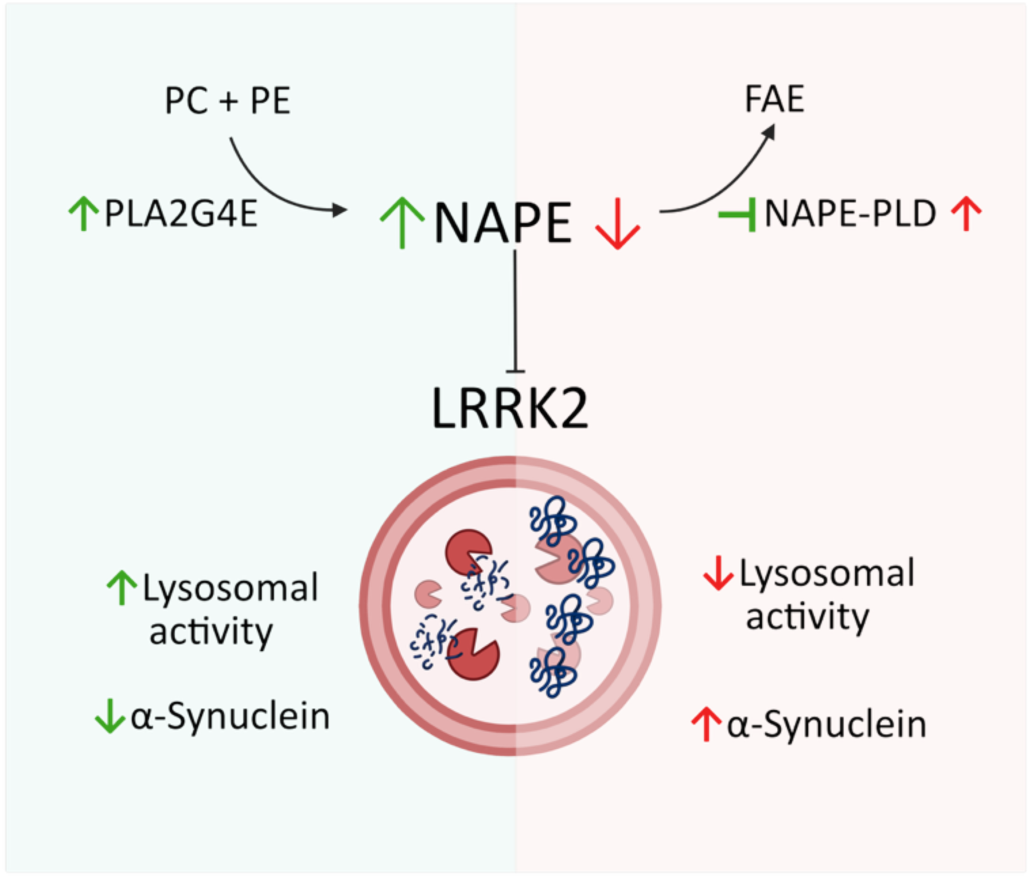
Schematic representation of the NAPE-LRRK2-Lysosome axis. NAPE accumulation, achieved either by enhancing their synthesis through PLA2G4E overexpression or by inhibiting their degradation through NAPE-PLD knockdown, decreases LRRK2 protein abundance and kinase activity. In turn, LRRK2 inhibitions enhances lysosomal activity, promoting α-synuclein clearance. Importantly, pharmacological inhibition of NAPE-PLD by LEI-401 rescues the lysosomal impairment induced by the pathogenic G2019S-LRRK2 variant.

LRRK2 is increasingly recognized as a stress-responsive regulator of lysosomes, recruited to enlarged or damaged lysosomes where it phosphorylates Rab GTPases and constrains lysosomal degradative activity (*36*, *50–52*). Importantly, other PD-linked proteins also shape this membrane-centered pathway: mutant VPS35 amplifies LRRK2 recruitment and Rab phosphorylation at lysosomes, thereby modulating endolysosomal homeostasis (*53*, *54*). Our data place NAPEs upstream of this pathway: elevating NAPEs by PLA2G4E overexpression or NAPE-PLD knockdown reduces LRRK2 abundance and Rab8a T72 phosphorylation and enhances lysosomal cathepsin B activity, whereas NAPE-PLD overexpression increases LRRK2 protein and kinase output and impairs lysosomal function. These manipulations support a model in which NAPE levels set a “lipid tone” that tunes LRRK2 activation and, consequently, lysosomal capacity (*35*, *36*, *38*). Mechanistically, this aligns with prior observations that NAPE accumulation reduces LRRK2 partitioning to membrane fractions, whereas enhanced NAPE hydrolysis promotes membrane association (*28*). NAPEs stabilize and remodel membranes (*23*, *24*), regulate fusion (*25*), and alter lipid raft organization (*26*), so changes in local NAPE content could influence LRRK2 recruitment to membrane microdomains supporting its activation. Unaltered TFEB responses in NAPE-enriched models suggest direct membrane effects, stabilizing fusion curvature or disrupting LRRK2-Rab raft recruitment. Our data do not yet discriminate whether NAPEs act directly on LRRK2 and its scaffolds or indirectly through altered Rab/VPS35 signaling and organelle organization; however, the tight correlation between NAPE levels, LRRK2 activity, and lysosomal performance supports a functionally integrated regulatory module. Additional studies will be required to fully define the molecular players involved in this NAPE-LRRK2-lysosome axis.

NAPE-PLD hydrolyzes NAPEs into bioactive FAEs such as OEA and PEA, as well as phosphatidic acid (PA), both of which can influence lysosomal function (*55*, *56*). FAEs can promote lysosomal biogenesis via PPAR-α–TFEB signaling (*57*, *58*),while PA modulates membrane properties in a concentration-dependent manner (*59*). However, our data argue that NAPEs themselves are the primary drivers of the observed effects. In NAPE-PLD-overexpressing cells, ionomycin-induced NAPE synthesis, which increases FAEs and PA, fails to restore lysosomal activity, whereas pharmacological NAPE accumulation via LEI-401 robustly enhances it. Although LRRK2 inhibition (MLi-2) partially rescues lysosomal defects, maximal restoration requires combined NAPE elevation, indicating both LRRK2-dependent and -independent contributions. Together, these findings support a model in which NAPEs act as stress-inducible membrane lipids with autonomous regulatory functions, effectively constraining LRRK2 activity and tuning lysosomal competence under proteotoxic conditions.

Pathogenic LRRK2 mutations, particularly G2019S, enhance kinase activity and drive lysosomal dysfunction, impaired autophagy, and α-synuclein accumulation (*38*, *47*). While LRRK2 kinase inhibitors can partially restore lysosomal function and reduce α-synuclein burden (*60*), our results extend this paradigm by identifying NAPEs as endogenous modulators of this axis. NAPE accumulation attenuates PFF-induced LRRK2 upregulation and enhances α-synuclein clearance in neuronal cells, whereas NAPE-PLD overexpression exacerbates both phenotypes. Importantly, in human iPSC-derived dopaminergic neurons carrying the G2019S mutation, NAPE-PLD inhibition restores lysosomal activity and reduces aggregate burden to levels comparable to isogenic controls. These findings integrate LRRK2 hyperactivation and α-synuclein pathology into a unified lipid-sensitive framework and suggest that modulation of NAPE metabolism can normalize lysosomal function even in a genetically sensitized context.

From a therapeutic perspective, NAPE-PLD inhibition emerges as a promising strategy to restore lysosomal proteostasis. By increasing endogenous NAPE levels, this approach enhances lysosomal function and promotes α-synuclein clearance while potentially avoiding some of the limitations associated with direct LRRK2 kinase inhibition. Small-molecule NAPE-PLD inhibitors effectively elevate NAPEs without compensatory increases in FAEs (*55*), supporting their utility as pharmacological tools. At the same time, given the broader roles of NAPEs and FAEs in brain physiology, including regulation of feeding, reward, and anxiety (*61*, *62*), careful in vivo studies will be required to define therapeutic windows and assess long-term effects. Nonetheless, targeting a stress-responsive lipid pathway offers a conceptually distinct and potentially complementary strategy to address convergent lysosomal defects observed in both genetic and sporadic PD (*60*).

While our study provides a robust framework for NAPE-mediated regulation of LRRK2 and lysosomal function, it has some limitations. First, most mechanistic experiments were performed in SH-SY5Y-derived neuronal cells, which only partially recapitulate mature nigral dopaminergic neuron biology. The rescue of lysosomal activity and α-synuclein handling in human G2019S dopaminergic neurons argues for disease relevance, but confirmation in in vivo (e.g., G2019S knock-in mice or α-synuclein PFF–injected rodents) will be important. Second, we focused on cathepsin B activity and aggregate counts; future studies should expand to broader lysosomal hydrolases, lysosomal pH, and TFEB/MiT–TFE transcriptional programs modulated by LRRK2 (*36*, *63*). Finally, the molecular interface between NAPEs and LRRK2 remains to be defined; which NAPE species are responsible, whether NAPEs alter Rab/scaffold recruitment (*35*, *50*, *52*), and how NAPE dynamics interact with other lipid pathways, including glucocerebrosidase-dependent sphingolipid metabolism. Addressing these questions will clarify how a stress-inducible membrane lipid precisely controls a disease-critical kinase and may reveal new opportunities to fine-tune lysosomal function in neurodegeneration.

## Methods

### Chemicals

MLi-2 LRRK2 inhibitor was purchased from Selleckchem (S9694). Bafilomycin A1 was purchased by Sigma-Aldrich (SML1661). Lei-401 NAPE-PLD inhibitor was kindly donated by Professor Van der Stelt (Leiden University), DOPE-N-Nonadecanoyl standard was purchased by Larodan (38–9091). Ionomycin was purchased by Sigma-Aldrich (I3909).

### Culture of Cell lines and treatments

Human neuroblastoma cell line SH-SY5Y (SK-N-SH-derived, hereafter “neuronal cells”) were cultured in RPMI1640 media (Euroclone, ECB2000L) and supplemented with 10% fetal calf serum (FCS) (Eurobio Scientific, CVFSVF00-01) and 1% penicillin-streptomycin (PS, Gibco, 15140-122; 100 units/mL final concentration). Cells were maintained at 37°C in a humidified CO^2^ incubator, and passaged using 0.05% Trypsin-EDTA (Gibco, 25300-054) when they reached 90% confluence. Cells were counted before each experiment, and seeded on autoclaved, 12 mm glass coverslips (uncoated) (Epredia, CB00120RA120MNZ0) for fixed-cell imaging, or on 35 mm glass-bottom microdishes (Ibidi GmbH, 80807-90) for live-cell imaging. Cells between passages 3 and 10 were used for experiments. MLi-2 LRRK2 inhibitor (Selleckchem) was dissolved in DMSO and diluted in full growth media to a final concentration of 10nM and incubated over-night. Bafilomycin A1 (Sigma-Aldrich, 500X) was dissolved in fresh full growth media and incubated for 4h prior to further analysis. LEI-401 was dissolved in in DMSO and diluted in full growth media to a final concentration of 1 μM and incubated over-night. Ionomycin was diluted in full growth media to a final concentration of 5 μM and incubated 30 minutes.

### Generation of NAPE-PLD knock-down cell lines

The NAPE-PLD Human Gene Knockout Kit (CRISPR, Origene, KN409877) was used following manufacturer’s instructions. Lipofectamine^TM^ 3000 Transfection Reagent (Thermo Fisher Scientific, L3000001) was used following standard protocol in OptiMEM media supplemented with 1% PS and 1% FCS. One day before transfection, SH-SY5Y cells were seeded in a six-well plate to reach 80-90% confluence at the time of transfection. Before transfection, culture medium was aspirated and 1715 µL of transfection medium was added. A mixture of donor DNA (10 μL per well), gRNA (20 µL per well), P3000 reagent (2 µL per well), and Lipofectamine 3000 (3 µL per well) was prepared in transfection medium (250 μl each) and incubated (15 min, room temperature). Transfection was performed by dropwise addition of the transfection mixture to the cells. Then, 6 h post-transfection, the culture medium was replenished with full growth media for additional 48 h. Cells were selected in the medium containing 1 μg/mL puromycin. Clonal cell lines were selected by clonal dilution. Three independent clones were used throughout this study.

### Generation of PLA2G4E-overexpressing cell lines

The pcDNA5/TO vector harboring EGFP-FL-PL2G4E was kindly donated by Professor Uyama (Kagawa University, (*33*)) and transfected into SH-SY5Y cells using 5 µg of DNA per well and Lipofectamine 3000 in OptiMEM media for 6h as described above. After 48 h from transfection, cells were selected with the medium containing 500 μg/mL Geneticin (Gibco, 10131-027). Clonal cell lines were generated by clonal dilution and maintained in the medium containing geneticin. Three of the clones expressing EGF-PLA2G4E were used throughout this study.

### Generation of NAPE-PLD-overexpressing cells

Lentiviral particles were produced by transient transfection of HEK293T cells using polyethyleneimine (PEI) (Sigma, #306185). Cells were seeded in 12-well plates at a density of 1.7 × 10⁵ cells per well and incubated for 24 hours to reach 70–80% confluency. Transfection mixtures were prepared in DMEM without FBS: the DNA mix contained 400 ng psPAX2 packaging plasmid (Addgene #12260), 350 ng VSV-G pseudotyping plasmid (Addgene #14888), 300 ng pRSV-Rev plasmid (Addgene #12253), and 650 ng NAPE-PLD-pLenti-C-mGFP-P2A-Puro transfer plasmid (OriGene #PS100093) (Fig. 1). A second mixture contained PEI diluted 1:50 in DMEM without FBS. The two mixtures were combined at a 1:1 ratio, vortexed gently, and incubated at room temperature for 20 minutes to allow PEI–DNA complex formation. Cell culture medium was replaced with fresh complete medium (DMEM + 10% FBS), and the transfection mixture was added dropwise to each well (total volume 100 µL per well). Cells were incubated at 37°C, 5% CO*₂* for 48 hours. Supernatants were then harvested, centrifuged at 3,000 rpm (1,100 × g) for 15 minutes at 4°C to remove cellular debris, and the clarified lentiviral particle-containing supernatant was aliquoted, snap-frozen in liquid nitrogen, and stored at −80°C. Viral titers were not formally quantified but estimated as sufficient for downstream transduction based on pilot GFP expression assays.

Stable SH-SY5Y NAPE-PLD-overexpressing cells were generated by transduction with the harvested lentiviral supernatant. SH-SY5Y cells were seeded in 12-well plates at 2 × 10⁵ cells per well and incubated for 24 hours. Medium was replaced with lentiviral supernatant supplemented with polybrene (Sigma, #H9268; final 8 µg/mL, 1:100 dilution from 0.8 mg/mL stock). Plates underwent spinoculation at 1,100 × g for 1 hour at 37°C to enhance transduction efficiency. Following spinoculation, plates were incubated for an additional 2 hours at 37°C, after which the medium was replaced with fresh DMEM/F-12 (1:1) supplemented with 2% FBS. Cells were recovered for 72 hours before initiating selection. Puromycin selection was performed in two sequential rounds: first at 5 µg/mL for 5 days, followed by 10 µg/mL for 7 days, with medium changes every 2–3 days. Transduction efficiency and selection success were confirmed by visualizing GFP fluorescence under a fluorescence microscope (Operetta CLS, Revvity, Milan). Selected polyclonal populations exhibited >95% GFP positivity and were expanded for downstream experiments.

### Cell culture experiments with hiPSC lines

Dopaminergic neurons were generated from neural progenitor cells (NPCs) according to an established protocol (*64*). Briefly, NPCs at 80% confluency were dissociated using Accutase for 5 minutes at 37 °C and collected via centrifugation (1100 rpm, 5 min). Cells were then seeded onto matrigel-coated 6-well plates (1:10 dilution in DMEM/F12) at a density of 1.0 × 10⁶ cells/well. Differentiation was initiated at day 0 by transitioning to differentiation medium. On day 6, the cells were passaged and re-seeded at 3.0 × 10⁶ cells/well. Neuronal maturation began on day 8 with the addition of 0.5 µM PMA, which was subsequently withdrawn on day 10. On day 14, the dopaminergic neurons were harvested and re-plated at a density of 2.6 × 10⁴ cells/cm, according to the experimental design planned. On day 21, cells were treated overnight with 1 μM LEI-401 and subsequently tested with magic red assay. Cells were treated with *α*-syn PFFs in the presence of vehicle or 1*μ*M LEI-401 for 48 h, followed by fixation and immunofluorescence analysis.

### Preparation of α-syn aggregates

*α*-Syn aggregates were prepared as described before (*65*). Briefly, human wild-type *α*-syn was purified from *Escherichia coli* BL21 (DE3) with RP-HPLC. Fibrils were then conjugated with Alexa Fluor™ 647 fluorophore (used in imaging experiments) (Invitrogen) using manufacturer’s labelling kit or not tagged with a fluorophore. Fibrils were kept at −80*°*C for long-term storage. Prior to exposure to cells, fibrils were diluted in growth medium at a working concentration of 500 nM and sonicated (BioBlock Scientific, Vibra Cell 7504) for 5 minutes at an amplitude of 80%, pulsed for 5 seconds “on” and 2 seconds “off”. Sonicated fibrils were then added on cells directly without the addition of any intracellular delivery agents for designated time points.

### Immunocytochemistry

Immunofluorescence on cells were performed using the standard protocol. Cells were fixed with 4% paraformaldehyde (PFA [Electron Microscopy Sciences, 15710]) for 20 minutes at room temperature (RT), followed by three washes with 1X DPBS. Cells were incubated in 1X DPBS-Tx (0.1% Triton-X100 [Sigma Aldrich, 9002-93-1] in 1X DPBS) for 3 minutes at RT, followed by incubation with blocking solution of 2% bovine serum albumin (BSA [Sigma Aldrich, A9647]) prepared in 1X DPBS for 1h at RT. Cells were then incubated with primary antibody overnight at 4*°*C. The following primary antibodies were used: rabbit anti-NAPE-PLD (Sigma Aldrich, HPA024338, 1:200), rabbit anti-PLA2G4E (Proteintech, 18088-1-AP, 1:400), rabbit anti-LRRK2 (Abcam, ab133474; 1:500) mouse anti-LAMP1 (Developmental Studies Hybridoma Bank, H4A3; 1:100), guinea pig anti-p62 (Progen, GP62-C; 1:500), rabbit anti-TFEB (Cell Signaling Technology, 4240; 1:100), chicken anti-MAP2 (BioLegend, 82250, 1:10,000), rabbit anti-TH (Pel-Freez, P40101, 1:1,000), and mouse anti-βIII-tubulin (BioLegend, 801202, 1:1,000). The next day, cells were washed three times with DPBS, followed by incubation with respective secondary antibodies (all from Thermo fisher Scientific, dilution of 1:500 in blocking solution) for 1h at RT. Cells were then washed three times with 1X DPBS, counterstained for nuclei with DAPI (1:1000 in PBS [Sigma-Aldrich, D9542]) and mounted on glass slides with Aqua-Poly/Mount (Polysciences Inc., 18606-20). Slides were imaged at least a day after mounting of coverslips.

### Magic red CTSB assay

Intracellular CTSB activity was measured in SH-SY5Y cells and dopaminergic iNeurons using the Magic Red™ CTSB assay kit from Abcam, ab270772, according to the manufacturer’s protocol. Cells were plated in glass-bottom matteks (Ibidi GmbH, 80807-90) the day before the experiment to reach a 70-80% confluence for neuronal cells, or two weeks before for iNeurons. Culture medium was gently removed and replaced with pre*-*warmed medium containing Magic Red substrate and Hoechst 33342 (Abcam, AB270772, 1:100). Ionomycin was diluted in the same media containing Magic Red to a final concentration of 1*μ*M. Cells were incubated at 37 °C, 5% CO*₂* for 30 min, protected from light. Following incubation, media was replaced with Phenol-free media and immediately live-imaged using a Nikon Eclipse Ti2 microscope with a spinning disk confocal set-up (Yokogawa) equipped with a live-cell incubation chamber that maintained the samples at 37 °C in a humidified atmosphere of 5% CO*₂* throughout imaging. Cells were imaged using a 100X oil immersion objective (1.45 numerical aperture). Optical sections of 0.5 *μ*m interval were taken to acquire the whole volume of the cells. Exposure time and gain were kept constant across all experimental conditions within each experiment. Analysis was performed by quantifying Magic Red fluorescence intensity in 40-60 randomly selected cells per experiment. A total of three independent experiments were conducted.

### Immunoblotting

Cell pellets were homogenized in 100 μl of radioimmunoprecipitation assay (RIPA) buffer consisting of 50 mM Tris-HCI (pH 7.4), 1% TritonX-100, 0.5% sodium-deoxycholate, 0.1% sodium dodecyl sulphate, 150 mM sodium chloride and 2 mM ethylenediaminetetraacetic acid, supplemented with 1X protease inhibitor (cOmplete Mini, EDTA-free; Sigma, 11836170001). Protein concentrations were measured using Bradford’s method, following manufacturer’s instructions (Thermo Fisher Scientific). Proteins (20 µg) were denatured in SDS (8%) and β-mercaptoethanol (5%) at 95 °C for 10 minutes. After separation by SDS-PAGE on a 4-20% gel (Biorad, 4568094), proteins were electro-transferred to nitrocellulose membranes. The membranes were blocked with Tropix I-Block buffer (Applied Biosystems, Thermo Fisher Scientific, T2015) and incubated overnight with primary antibodies (anti-NAPE-PLD, Sigma-Aldrich, HPA024338, 1:500; anti-PLA2G4E, Proteintech, 18088-1-AP, 1:1000; anti Rab8a, Abcam, Ab188574, 1:500; anti-pT72Rab8a, Abcam, Ab230260, 1:1000; anti-LC3, Cell Signaling Technologies, #3868, 1:1000; anti-GAPDH antibody Sigma-Aldrich, G9545, 1:5000; anti-Actin, MP Biomedicals, 691001, 1:1000) in blocking buffer, followed by incubation with horseradish peroxidase-conjugated secondary antibody (Millipore, 1:5000) in TBS containing 0.1% Tween-20 at room temperature for 1 h. Finally, proteins were visualized using an ECL kit (Thermo Fisher Scientific, 34580) and chemiluminescence images were acquired using GE Amersham Imager AI680 analyzer.

### mRNA isolation and qPCR

Total RNA was prepared from cell pellets using the Ambion PureLink RNA minikit (Invitrogen, Thermo Fisher Scientific, 12183018A) as directed by the supplier. Samples were treated with DNase (PureLink DNase, Invitrogen, Thermo Fisher Scientific, 12185010) and cDNA synthesis was carried out using the high-capacity cDNA reverse transcription kit (Applied Biosystems, Thermo Fisher Scientific 4368814) according to the manufacturer’s protocol using purified RNA (0.5-1 μg). First-strand cDNA was amplified using the iTaq universal SYBR Green supermix (Bio-Rad, 1725124). Primers sequences are indicated in **Table 1**. Copy numbers of cDNA targets were quantified at the cycle threshold. Quantitative PCR was performed in 384-well PCR plates and run at 95°C for 10 min, followed by 40 cycles, each cycle consisting of 15 s at 95°C and 1 min at 60°C, using a CFX 384-Real Time System (Bio-Rad). The BestKeeper software (*66*) was used to determine expression stability and geometric mean of three different housekeeping genes (Actin, GAPDH and HPRT). ΔCt values were calculated by subtracting the Ct value of the geometric mean of these housekeeping genes from the Ct value for the genes of interest. The relative quantity of genes of interest was calculated by the expression 2−ΔΔCt. Results are reported as fold induction over control.

**Table 1:**
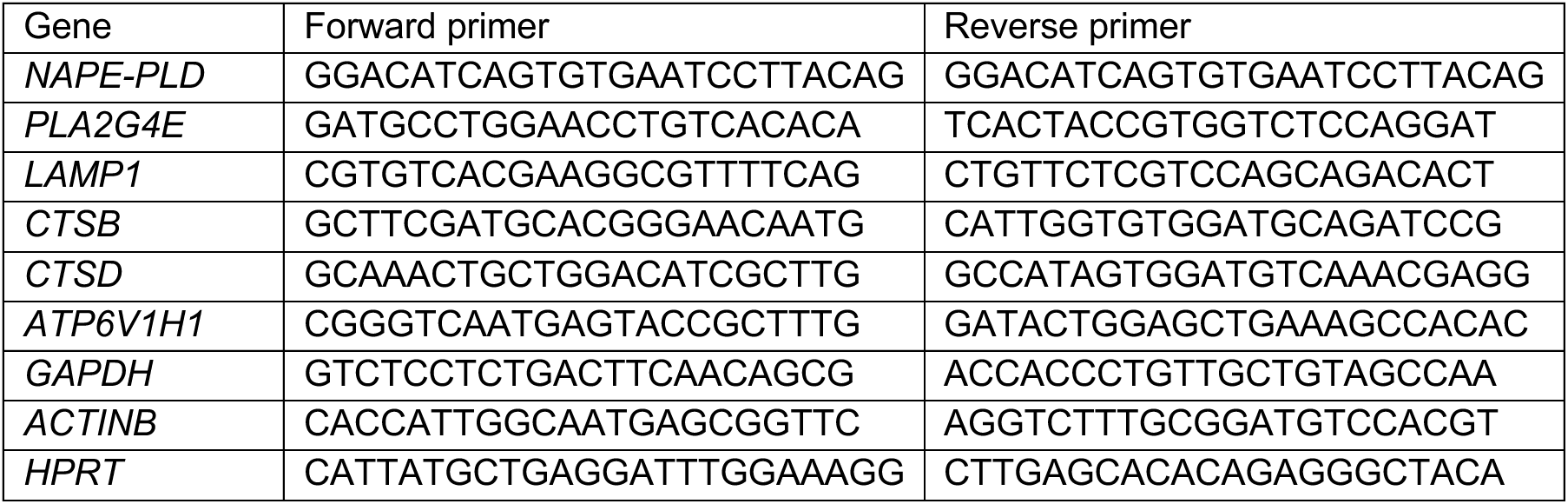
Primers used in the study.

### Lipid extraction and LC/MS analyses

NAPE levels were measured as previously described (*29*). Briefly, snap-frozen cell pellets were homogenized in a chloroform:methanol mixture (1:2 v/v, 2 mL) containing DOPE-N-Nonadecanoyl (Larodan, #38-9091) as internal standards. Lipids were extracted twice with chloroform (0.6 mL) and the organic phases were washed with water (0.6 mL), collected, and dried under nitrogen stream. The organic extracts from cells were reconstituted in a mix of methanol/chloroform (9:1 v/v, 0.1 mL). Targeted analysis of the samples was carried out on a Waters Acquity UPLC system coupled with a Xevo TQ-MS triple quadrupole mass spectrometer. NAPEs were separated on a reversed-phase BEH C18 column (2.1 x 50 mm, 1.7 μm), whose temperature was kept at 55°C, and eluted at a flow rate of 0.450 mL/min using the following gradient conditions: A = 10 mM ammonium formate in acetonitrile/water (60:40 v/v), B = 10 mM ammonium formate in isopropyl alcohol/acetonitrile (90:10 v/v); 0.0-1.0 min 40% B, 1.0-7.0 min 40 to 100% B, 7.0-8.0 min at 100% B. The column was then reconditioned at 40% B for 1 min. The total run time was 9 min and the injection volume was set to 5 µL. The mass spectrometer operated in the positive ESI mode and analytes were quantified by multiple reaction monitoring (MRM), following the transitions reported in Table 2. The capillary and the cone voltages were set at 3 kV and 30 V for all transitions, respectively. The source temperature was set to 120 °C. Desolvation and cone gas flows (N_2_) were set to 800 and 50 l/h, respectively. Desolvation temperature was set at 450°C. Data were acquired by MassLynx software and quantified by TargetLynx software. Quantification of NAPE species was performed by means of a six-point calibration curve where standards were prepared by spiking the only non-endogenous synthetic NAPE reference analytical standard we could purchase, DOPE-N-19 (18:1/18:1/N19:0) in matrix at six different concentrations, from 1 to 500 nM. As this LC/MS-MS protocol does not allow one to differentiate sn-1 from sn-2 substituents, individual NAPE species are designated below as NAPE (X:Y-N-acyl, **Table 2**), where X is the total number of carbon atoms and Y the total number of double bonds in the sn-1 and sn-2 chains.

**Table 2:** Parent, quantifier and qualifier ions (as m/z) tracked for each of the 14 NAPEs under investigation. The corresponding observed retention time (in minutes) is also indicated.

| ID | NAME | Parent m/z | Fragment m/z<br>(Qn) | Fragment m/z<br>(Qf) | Rt (min) |
| --- | --- | --- | --- | --- | --- |
| 1 | PE (P50:1-N-acyl) | 940.8 | 282.3 | 561.5 | 5.03 |
| 2 | PE (P52:2-N-acyl) | 966.8 | 282.3 | 587.5 | 5.03 |
| 4 | PE (P54:5-N-acyl) | 988.8 | 282.3 | 609.5 | 4.79 |
| 5 | PE (P54:4-N-acyl) | 990.8 | 282.3 | 611.5 | 5.00 |
| 7 | PE (P50:6-N-acyl) | 1014.8 | 308.3 | 609.5 | 4.81 |
| 10 | PE (P50:5-N-acyl) | 1016.8 | 308.3 | 611.5 | 5.01 |
| 11 | PE (P50:5-N-acyl) | 1016.8 | 310.3 | 609.5 | 5.00 |
| 20 | PE (P50:6-N-acyl) | 1014.8 | 282.3 | 635.5 | 4.92 |
| 21 | PE (P50:6-N-acyl) | 1014.8 | 310.3 | 607.5 | 4.92 |
| 22 | PE (50:6-N-acyl) | 1030.8 | 282.3 | 651.5 | 4.84 |
| 23 | PE (50:6-N-acyl) | 1030.8 | 310.3 | 623.5 | 4.82 |
| 24 | PE (50:4-N-acyl) | 1034.8 | 310.3 | 627.5 | 5.10 |
| 25 | PE (P58:6-N-acyl) | 1042.8 | 310.3 | 635.5 | 4.98 |
| 26 | PE (58:6-N-acyl) | 1058.8 | 310.3 | 651.5 | 5.03 |

### Fixed-cell microscopy

Images were acquired using a Nikon Eclipse Ti2 microscope with a spinning disk confocal set-up (Yokogawa) equipped with four lasers (wavelength in nm): 405, 488, 561 and 640 nm. Samples were imaged using a 100X oil immersion objective (1.45 numerical aperture) with a 1x zoom. The entire cell volume was imaged for all samples, with optical sections of 0.5 *μ*m.

SIM was performed on a Zeiss Elyra 7 Lattice SIM microscope (Carl Zeiss, Germany) using C Plan-Apochromat 63X/1.46 oil objective with a 1.518 refractive index oil (Carl Zeiss). The fluorescence signal was detected on PCO Edge 4.2 QE 82% cameras. Raw images were composed of thirteen images per plane per channel and acquired with a Z-distance of 251 nm. Cameras were aligned using an internal pattern and tetraspeck beads of 100 nm were used to correct chromatic aberration.

### Quantification and Statistical Analyses

Images were processed for analysis using FIJI software (). Colocalization analysis was performed using the JACoP plug-in FIJI (*67*). 3D images were used for colocalizing two channels with object-based detection, and thresholding of objects was performed manually.

No prior power analysis was done to measure the sample size. Graphs were plotted using GraphPad prism 10.0, and appropriate statistical tests were performed on raw data. For all datasets, an initial normality distribution test was performed, and non-parametric tests were performed for any datasets that did not satisfy normal distribution. Statistical tests performed are mentioned in the figure legends. All the experiments were performed for three independent biological replicates.

## Acknowledgments

We are grateful to Prof. Toro Uyama and Prof. Natsuo Ueda for generously providing the EGFP-FL-PL2G4E construct, and to Prof. Mario van der Stelt for providing the NAPE-PLD inhibitor LEI-401. We thank Prof. Takashi Nonaka for providing the α-synuclein fibrils. We are grateful to Dr. Valeria Valente and Dr. Ranabir Chakraborty for critical reading of the manuscript; we also thank Dr. Ranabir Chakraborty for assistance with super-resolution image processing. We thank Dr. Audrey Salles for technical assistance with super-resolution microscopy and the kind financial support for the use of the Zeiss Elyra 7 microscope, Institut Pasteur, France–BioImaging infrastructure network (FBI) supported by the French National Research Agency (ANR-10-INBS-04; Investments for the Future), the Région Ile-de-France (program DIM1HEALTH), and the French Government Investissement d’Avenir Programme-Laboratoire d’Excellence ‘Integrative Biology of Emerging Infectious Diseases’ (ANR-10-LABX-62-IBEID). We also thank all members of the Membrane Traffic and Pathogenesis Unit at the Institut Pasteur for insightful discussions. Finally, we thank Reine Bouyssie, from the administrative staff of the Membrane Traffic and Pathogenesis Unit, for her continued support.

The present study was funded by MSCA-IF-Fellowship LipiSyn – 101064077 (FP) and Fondation Alzheimer Young Researcher Program, 990987 (FP), France Parkinson - Soutien de l’Association France Parkinson AAP 2021 (CZ), Don Explore AD - Programme Explore de l’Institut Pasteur (CZ), Agence Nationale de la Recherche ANR-20-CE13-0032-01 (CZ), ANR-23-CE16-0012-03 UnProSec (CZ), DFG-ANR #490761034 (MD), Fondation pour la Recherche Médicale FRM - EQU202103012692 (CZ), FRMMND202310017892 (CZ), Fondation Alzheimer AAP 2024 (CZ), ERC CoG – 101003329 (MD).

## Author Contributions

Conceptualization: FP; Methodology: FP, CG, SS, GO, GS; Investigation: FP; Visualization: FP; Supervision: FP, ML, MD, AA; Funding acquisition: FP, CZ; Writing – original draft: FP; Writing – review & editing: FP, CG, MD, CZ.

## Data Availability

All data needed to evaluate the conclusions in the paper are present in the paper and/or the Supplementary Materials.

