## supplementary figures for "An *N*-acylphosphatidylethanolamine-LRRK2 axis controls lysosomal homeostasis in Parkinson’s disease"

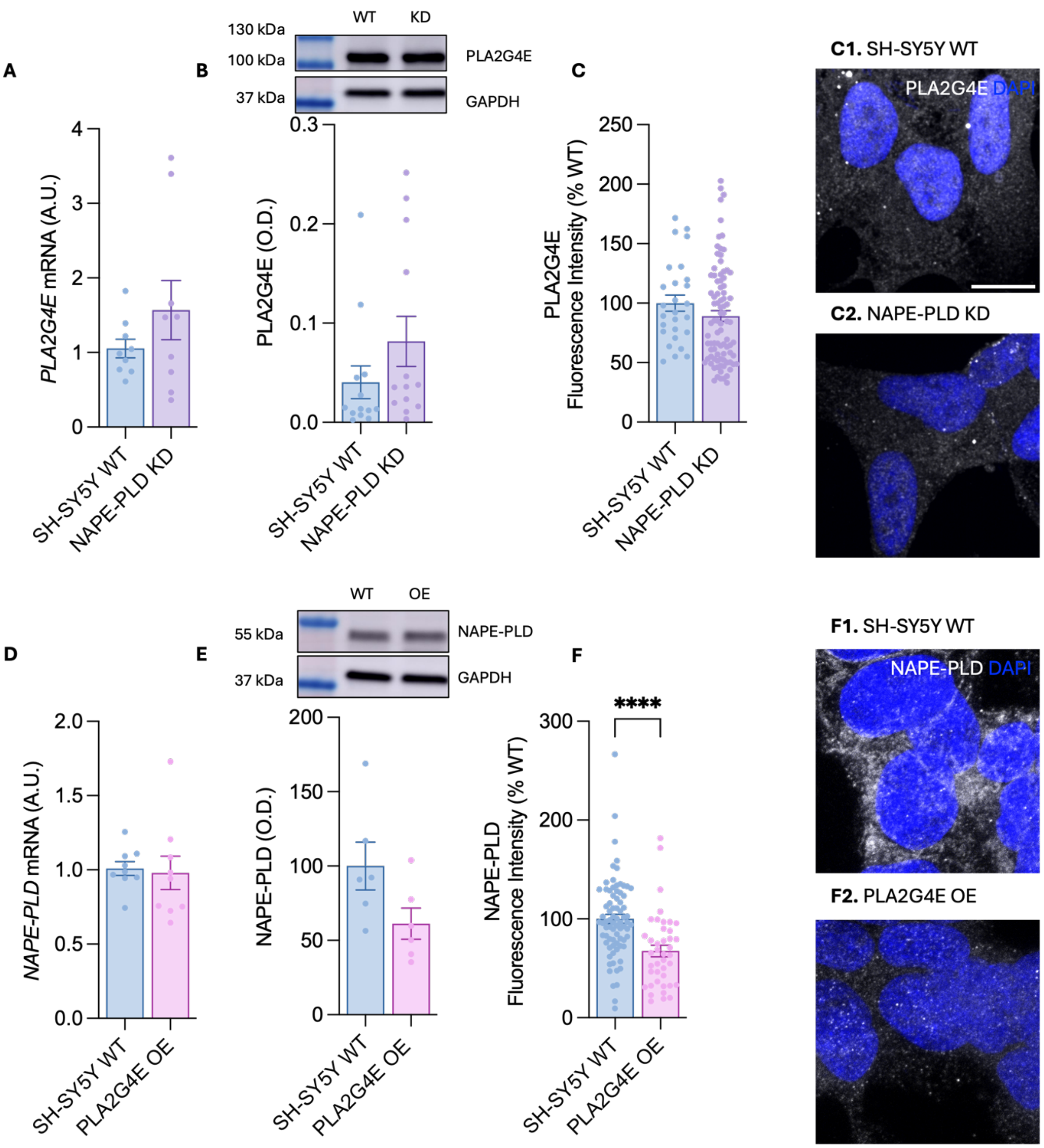


**Supplementary Figure S1. Characterization of NAPE metabolism in catecholaminergic SH-SY5Y neurons accumulating NAPEs.** (**A**) *PLA2G4E* mRNA levels in NAPE-PLD-knockdown (KD, violet bars) and wild-type (WT, blue bars) cells. (**B**) Western blot analysis of PLA2G4E protein levels in KD and WT cells. Top, representative blot; bottom, densitometric quantification normalized to GAPDH. (**C**) Immunofluorescence analysis of PLA2G4E (gray) with representative images (**C1, C2**) and quantification. Nuclei were counterstained with DAPI. (**D**) *NAPE-PLD* mRNA levels in PLA2G4E-overexpressing (OE, pink bars) and WT (blue) cells. (**E**) Western blot analysis of NAPE-PLD protein levels in OE and WT cells. Top, representative blot; bottom, densitometric quantification normalized to GAPDH. (**F**) Immunofluorescence analysis of NAPE-PLD (gray) with representative images (**F1, F2**) and quantification. Nuclei were counterstained with DAPI. Error bars represent SEM. ****P < 0.0001 Student's *t*-test with Welch's correction. Scale bars, 10 μm.

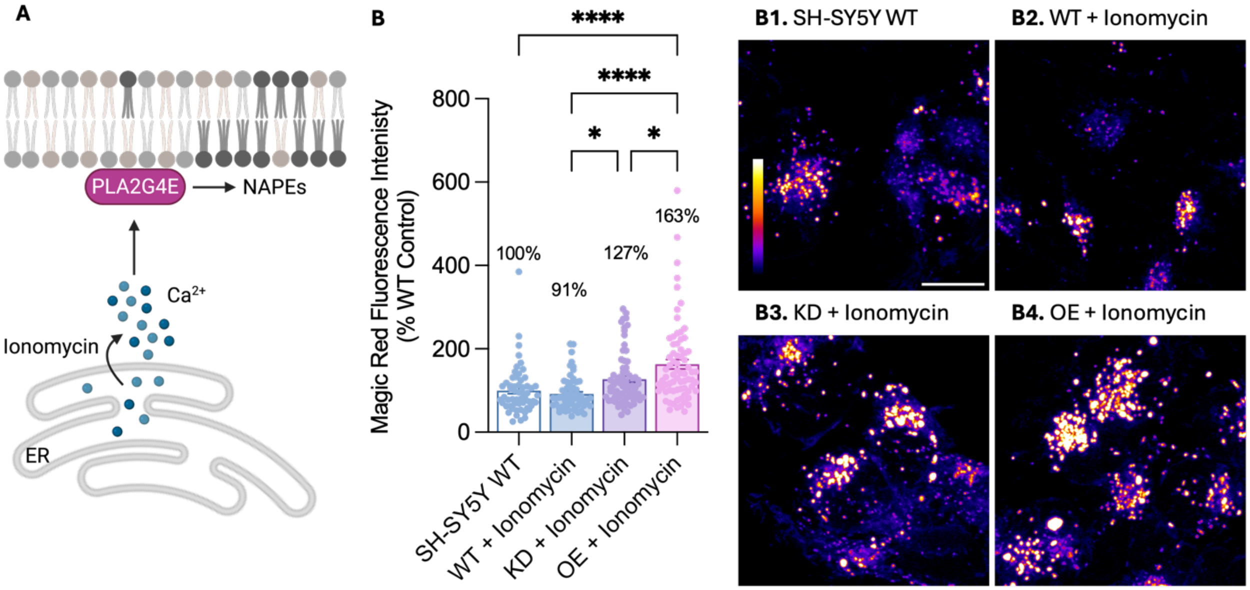


**Supplementary Figure S2. Stimulation of NAPE accumulation further enhances lysosomal activity in neuronal cells.** (**A**) Schematic representation of the experimental strategy. Ionomycin treatment induces intracellular $Ca^{2+}$ release, promoting the production and accumulation of NAPEs in PLA2G4E-overexpressing (OE) and NAPE-PLD knockdown (KD) cells, respectively. Ionomycin was administered together with the Magic Red reagent for 30 min before image acquisition. (**B**) Fluorescence intensity quantification and representative images of the Magic Red – cathepsin B assay in untreated WT cells (**B1**) and WT (**B2**), NAPE-PLD KD (**B3**), and PLA2G4E OE (**B4**) cells treated with ionomycin. Representative images are displayed using the FIRE lookup table (LUT) to indicate fluorescence intensity. Scale bars, 10 μm. Mean values are indicated in the graphs; error bars represent SEM. *P < 0.05, ****P < 0.0001; one-way ANOVA with Tuckey's multiple-comparisons test.

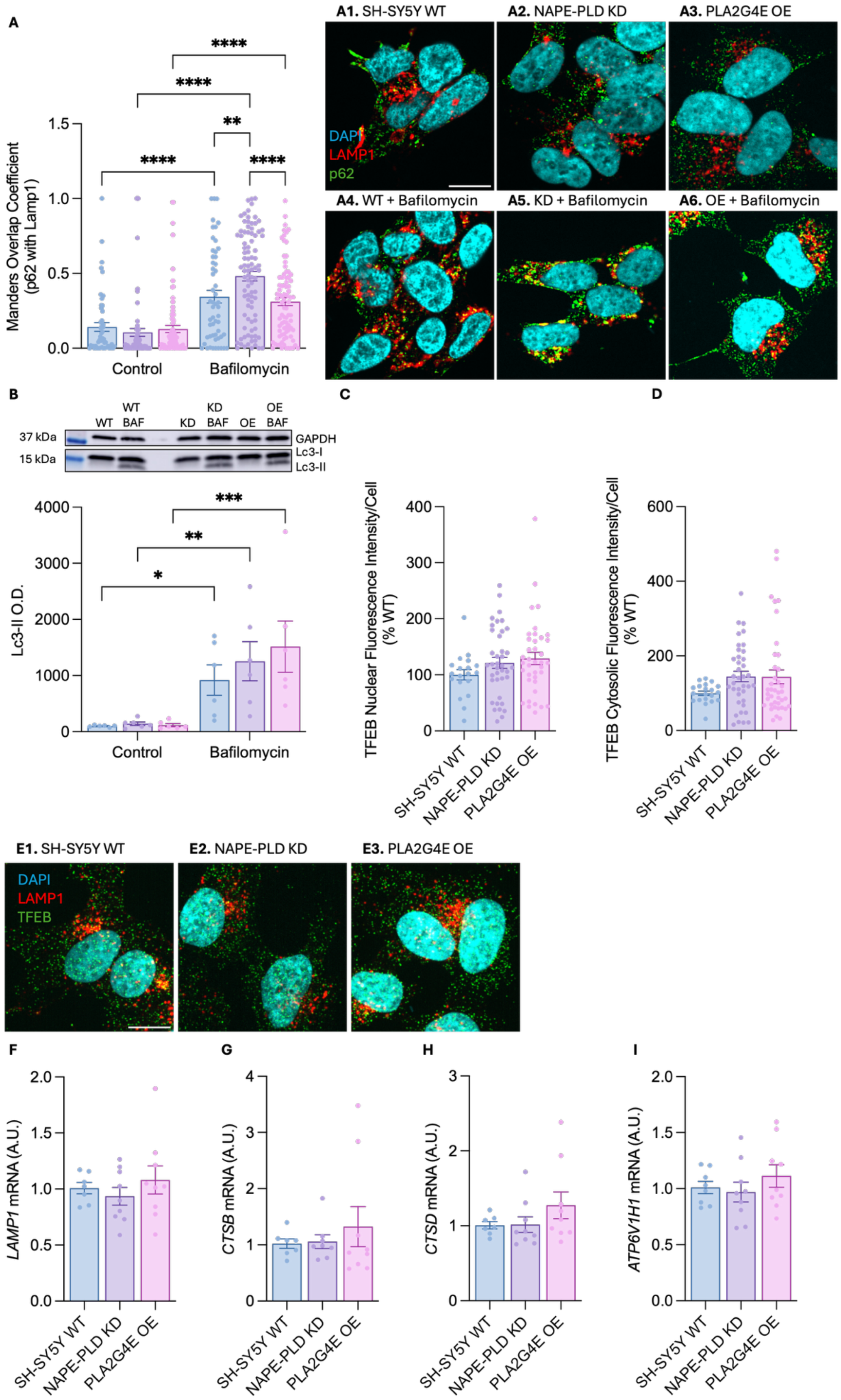

A
A1. SH-SY5Y WT
A2. NAPE-PLD KD
A3. PLA2G4E OE
A4. WT + Bafilomycin
A5. KD + Bafilomycin
A6. OE + Bafilomycin
DAPI
LAMP1
p62
Manders Overlap Coefficient (p62 with Lamp1)
Control
Bafilomycin
B
WT
WT BAF
KD
KD BAF
OE
OE BAF
37 kDa
15 kDa
GAPDH
Lc3-I
Lc3-II
Lc3-II O.D.
C
TFEB Nuclear Fluorescence Intensity/Cell (% WT)
D
TFEB Cytosolic Fluorescence Intensity/Cell (% WT)
SH-SY5Y WT
NAPE-PLD KD
PLA2G4E OE
E1. SH-SY5Y WT
E2. NAPE-PLD KD
E3. PLA2G4E OE
TFEB
F
LAMP1 mRNA (A.U.)
G
CTSB mRNA (A.U.)
H
CTSD mRNA (A.U.)
I
ATP6V1H1 mRNA (A.U.)

**Supplementary Figure S3. Autophagic flux is preserved in NAPE-enriched cells.** (**A**) Quantification and representative images of the colocalization between LAMP1-positive structures and p62 in WT (**A1, A4**), NAPE-PLD KD (**A2, A5**) and PLA2G4E OE (**A3, A6**) cells treated or not with Bafilomycin A1, to inhibit autophagic flux. Colocalization is expressed as Mander's overlap coefficient for the fraction of p62 signal overlapping with LAMP1. LAMP1 is shown in red, p62 in green, and nuclei were counterstained with DAPI. (**B**) Western blot analysis of LC3-II protein levels in WT, NAPE-PLD KD and PLA2G4E OE cells treated or not with Bafilomycin A1. Top, representative blot; bottom, densitometric quantification normalized to GAPDH. (**C-E**) Immunofluorescence analysis of TFEB (green) and LAMP1 (red) with representative images and quantification in WT (**D1**; blue), NAPE-PLD KD (**D2**; violet), and PLA2G4E OE (**D3**, pink) cells. Nuclei were counterstained with DAPI. Scale bar, 10 μm. (**F-I**) *LAMP1* (**F**), *CTSB* (**G**), *CTSD* (**H**), *and ATP6V1H1* (**I**) mRNA levels in NAPE-PLD KD, PLA2G4E OE and WT cells under basal conditions. Error bars represent SEM. **P < 0.01, ****P < 0.0001; two-way ANOVA with Tuckey's multiple-comparisons test.

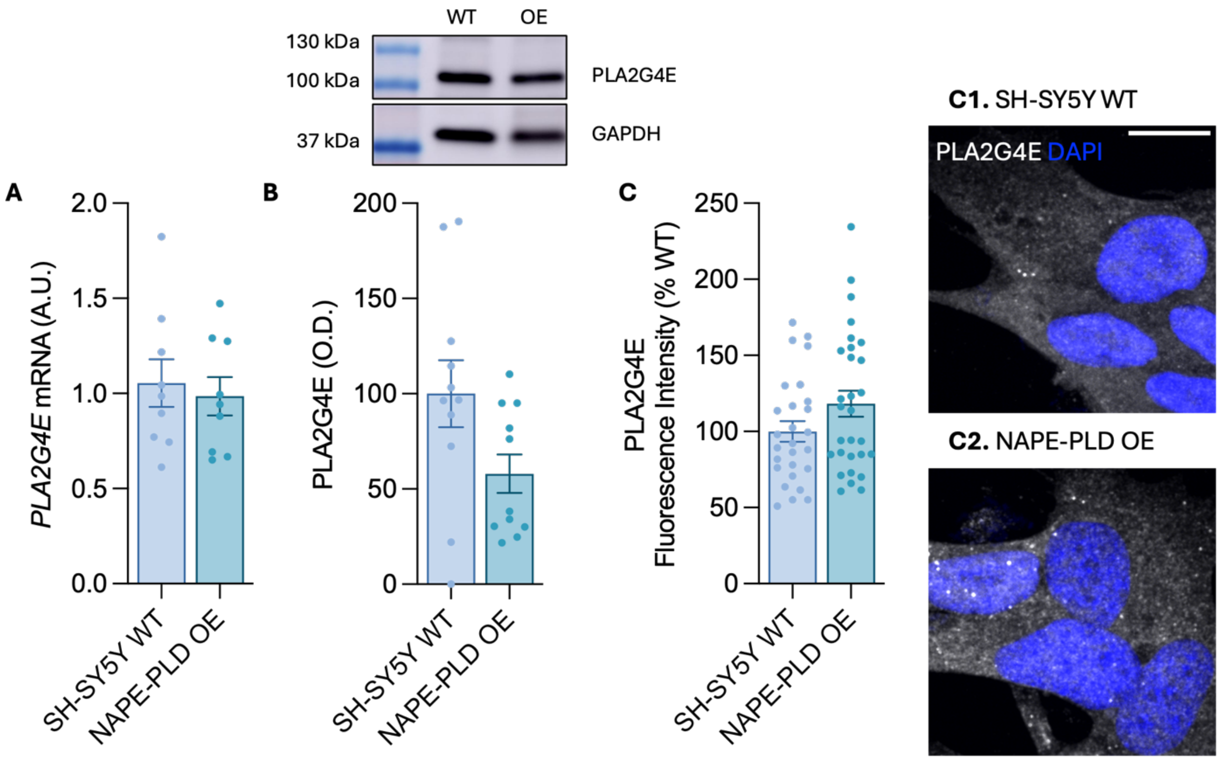


**Supplementary Figure S4. Characterization of PLA2G4E in NAPE-PLD-overexpressing SH-SY5Y neuronal cells.** (**A**) *PLA2G4E* mRNA levels in NAPE-PLD-overexpressing (OE; green) and wild-type (WT; blue) cells. (**B**) Western blot analysis of PLA2G4E protein levels in OE and WT cells. Top, representative blot; bottom, densitometric quantification normalized to GAPDH. (**C**) Immunofluorescence analysis of PLA2G4E (gray) with representative images (**C1, C2**) and quantification. Nuclei were counterstained with DAPI (blue). Scale bar: 10 μm. Error bars represent SEM. Student's *t*-test with Welch's correction.

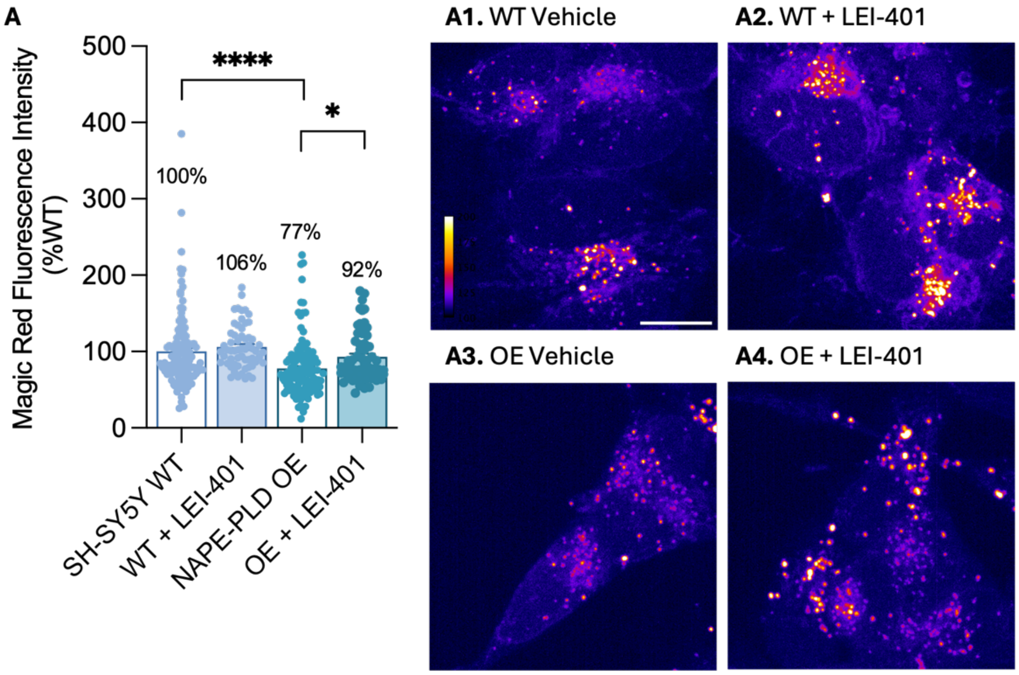


**Supplementary Figure S5. Pharmacological inhibition of NAPE-PLD restores lysosomal activity in NAPE-PLD overexpressing SH-SY5Y cells.** (**A**) Fluorescence intensity quantification and representative images of the Magic Red – cathepsin B assay, indicating lysosomal degradative capacity per cell, in WT cells (**A1, A2**) and NAPE-PLD OE cells (**A3, A4**) cells treated with vehicle (**A1, A3**) or the NAPE-PLD inhibitor LEI-401 (**A2, A4**). Representative images are displayed using the FIRE lookup table (LUT) to indicate fluorescence intensity. Scale bar, 10 μm. Mean values are indicated in the graphs; error bars represent SEM. *P < 0.05, ****P < 0.0001; two-way ANOVA with Tuckey's multiple-comparisons test.

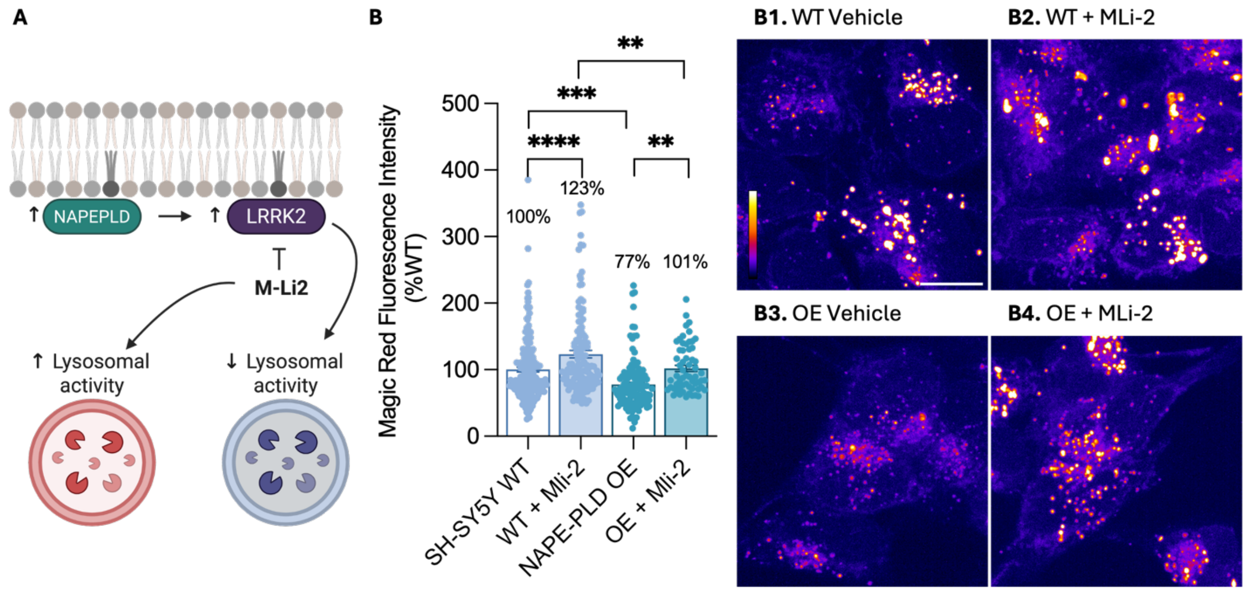


**Supplementary Figure S6. Pharmacological inhibition of LRRK2 enhances lysosomal activity in wild-type and NAPE-PLD overexpressing SH-SY5Y cells.** (**A**) Schematic representation of the experimental strategy. NAPE-PLD overexpressing cells exhibit increased LRRK2 protein expression and activation. The effect of LRRK2 inhibition on lysosomal activity was assessed by overnight treatment with MLi-2 followed by the Magic Red – cathepsin B assay. (**B**) Fluorescence intensity quantification and representative images of the Magic Red – cathepsin B assay in WT (**B1, B2**) or NAPE-PLD OE (**B3, B4**) cells treated with vehicle (**B1, B3**) or MLi-2 (**B2, B4**). Representative images are displayed using the FIRE lookup table (LUT) to indicate fluorescence intensity. Scale bar, 10 μm. Mean values are indicated in the graphs; error bars represent SEM. **P < 0.01, ***P < 0.001, ****P < 0.0001; two-way ANOVA with Tuckey's multiple-comparisons test.

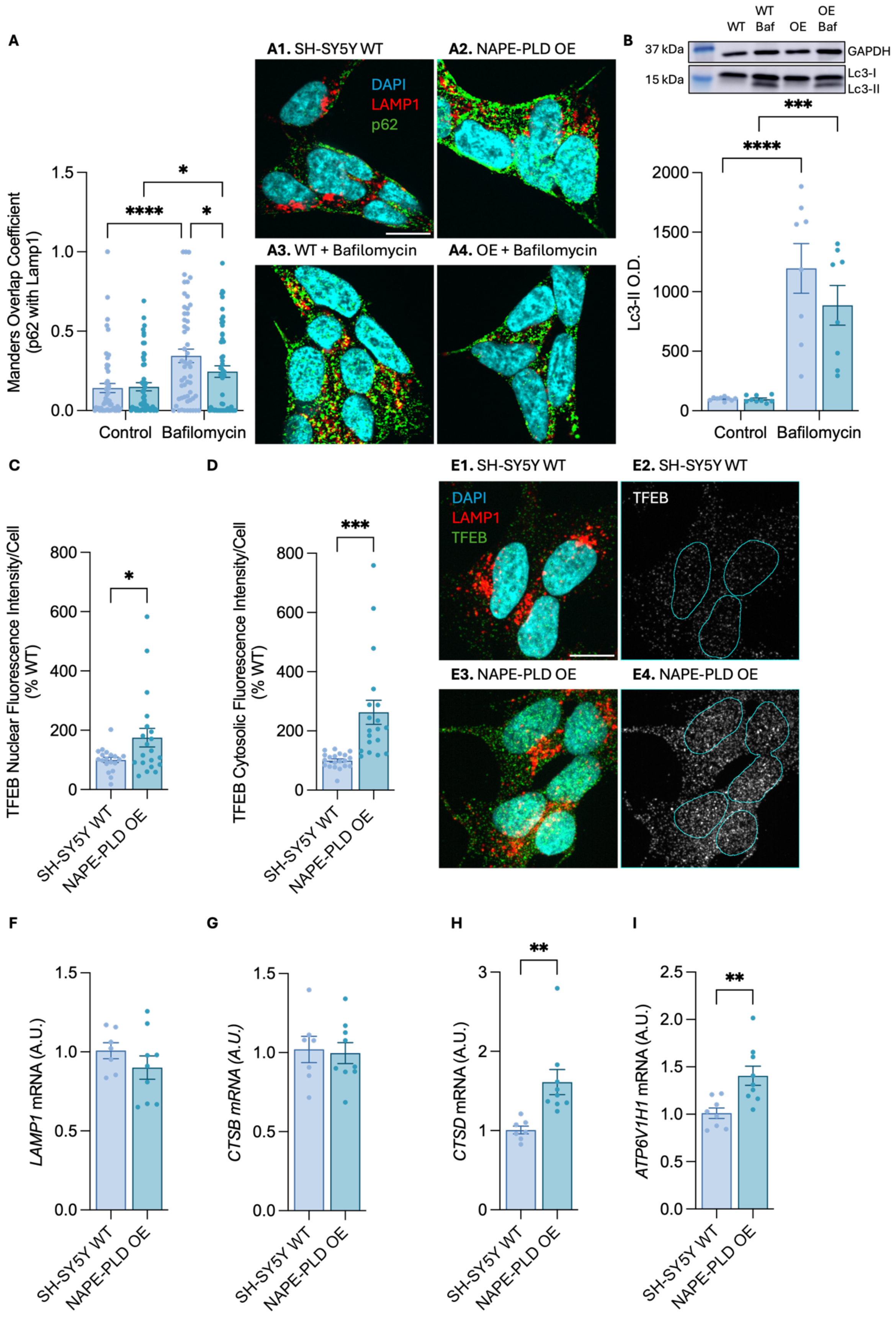

A
A1. SH-SY5Y WT
A2. NAPE-PLD OE
DAPI
LAMP1
p62
A3. WT + Bafilomycin
A4. OE + Bafilomycin
Manders Overlap Coefficient (p62 with Lamp1)
Control
Bafilomycin
B
WT
WT Baf
OE
OE Baf
37 kDa
GAPDH
15 kDa
Lc3-I
Lc3-II
Lc3-II O.D.
Control
Bafilomycin
C
TFEB Nuclear Fluorescence Intensity/Cell (% WT)
SH-SY5Y WT
NAPE-PLD OE
D
TFEB Cytosolic Fluorescence Intensity/Cell (% WT)
SH-SY5Y WT
NAPE-PLD OE
E1. SH-SY5Y WT
E2. SH-SY5Y WT
DAPI
LAMP1
TFEB
TFEB
E3. NAPE-PLD OE
E4. NAPE-PLD OE
F
LAMP1 mRNA (A.U.)
SH-SY5Y WT
NAPE-PLD OE
G
CTSB mRNA (A.U.)
SH-SY5Y WT
NAPE-PLD OE
H
CTSD mRNA (A.U.)
SH-SY5Y WT
NAPE-PLD OE
I
ATP6V1H1 mRNA (A.U.)
SH-SY5Y WT
NAPE-PLD OE

**Supplementary Figure S7. NAPE-PLD overexpression alters autophagic flux in SH-SY5Y cells.** (**A**) Quantification and representative images of the colocalization between LAMP1-positive structures and p62 in WT (**A1, A3**) and NAPE-PLD OE (**A2, A4**) cells treated or not with bafilomycin A1 to inhibit autophagic flux. Colocalization is expressed as Mander's overlap coefficient for the fraction of p62 signal overlapping with LAMP1. LAMP1 is shown in red, p62 in green, and nuclei were counterstained with DAPI. Error bars represent SEM. *P < 0.05, ****P < 0.0001; two-way ANOVA with Tuckey's multiple-comparisons test. (**B**) Western blot analysis of LC3-II protein levels in WT, and NAPE-PLD OE cells treated or not with Bafilomycin A1. Top, representative blot; bottom, densitometric quantification normalized to GAPDH. (**C-E**) Immunofluorescence analysis of TFEB (green) and Lamp1 (red) with representative images and quantification in WT (**E1, E2**) and NAPE-PLD OE (**E3, E4**) cells. TFEB localization was quantified in the nuclear (**C**) and cytosolic (**D**) compartments. Nuclei were counterstained with DAPI (cyan). Scale bar, 10 μm. (**F-I**) *LAMP1* (**F**), *CTSB* (**G**), *CTSD* (**H**), *and ATP6V1H1* (**I**) mRNA levels in WT and NAPE-PLD OE cells under basal conditions. Error bars represent SEM. *P < 0.05, ***P < 0.001, ****P < 0.0001; two-way ANOVA with Tukey's multiple-comparison test (**A**) and Student's t test with Welch's correction (**B–D**).

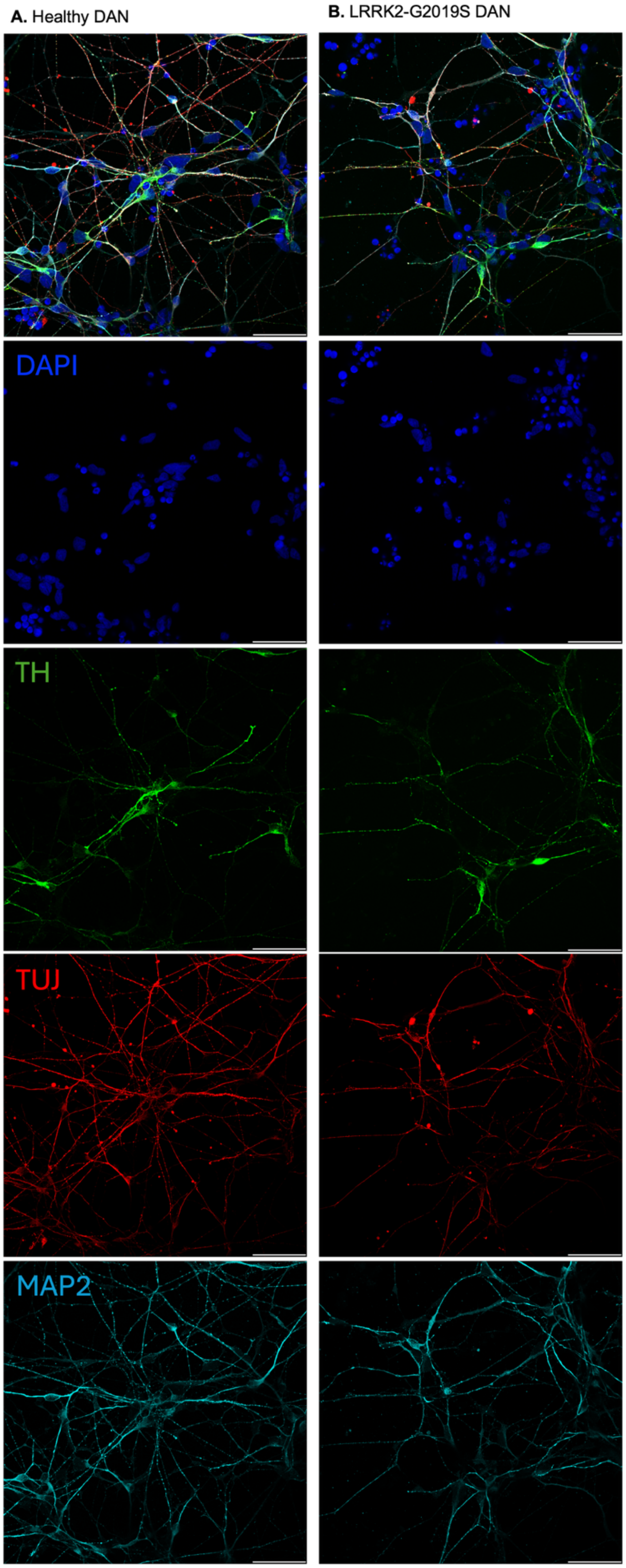
A. Healthy DAN
B. LRRK2-G2019S DAN
DAPI
TH
TUJ
MAP2

**Supplementary Figure S8. Characterization of human iPSC-derived dopaminergic neurons.** (**A, B**) Representative immunofluorescence images of healthy (**A**) and G2019S-LRRK2 mutant dopaminergic neurons (DANs; **B**) stained for tyrosine hydroxylase (TH; green), neuron-specific βIII-tubulin (TUJ; red) and microtubule-associated protein 2 (MAP2; cyan). Nuclei were counterstained with DAPI (blue). Scale bar, 50 μm.
